# Isolation of antibiotic Producing Microorganisms from some water bodies within Eastern and Greater Accra Regions of Ghana

**DOI:** 10.1101/695783

**Authors:** Daniel Amiteye, Nicholas Tete Kwaku Dzifa Dayie, Stephen Yao Gbedema, Vivian Etsiapa Boamah, Francis Adu, Marcel Tunkumgnen Bayor

## Abstract

**Background:** Most antimicrobial agents used today are isolated and extracted from microbial source. The emergence of antimicrobial resistance and need for better, broad spectrum antimicrobial agent is always in high demand. In this study, a total of 112 aquatic microbial isolates from 14 sample sites of water bodies in Ghana were screened by agar-well diffusion method for the ability of antimicrobial metabolites.

**Results:** Out of these antibacterial activities, 10 inhibited the growth of at least one test microorganism with zones of growth inhibitions ranged between 2.5±0.5 - 35.5±0.5 mm against *Staphylococcus aureus* (ATCC25923), *Streptococcus pyogenes* (clinical isolates)*, Escherichia coli* (NCTC9002), *Pseudomonas aeruginosa* (ATCC27853), *Neisseria gonorrhoeae* (clinical isolate), *Klebsiella pneumonia*e (Clinical isolate), *Bacillus subtilis* (NCTC10073) and *Salmonella typhi* (NCTC 6017). The extracts of the isolates showed minimum inhibitory concentrations of which extract NKSEW_3_ against *Escherichia coli, Klebsiella pneumoniae* and *Pseudomonas aeruginosa* were 25.00, 12.50 and 25.00 mg/mL respectively while that of NKLS_6_ were 12.50, 6.25 and 25.00 mg/mL. The isolates NKSEW_3_ and NKLS_6_ were identified to be *Pseudomonas species* using chromagar and biochemical tests. The GC-MS result for NKLS_6_2 (a fraction obtained from NKLS_6_) revealed five compounds including; Tridecanal, 17-octadecanal, ethanol, 2-(9-octadecenyloxy)-, (Z), 2-pentadecanone, 6, 10, 14-trimethyl diisooctyl phthalate and 17-octadecanal (C_18_H_34_O) as good pharmacological agents.

**Conclusion:** Microorganisms isolated from water bodies in Ghana have the potential of producing antimicrobial agents.

**Author Summary:** In this study we use the agar well-diffusion to screen the isolates of water and soil samples collected within Greater Accra and Eastern Regions in Ghana against the test organisms.

## Background

The problem of resistance against the present antimicrobial agents in pathogens increases daily. Hence, new and effective antimicrobial agents are needed to combat these foes^1^. The sources of antimicrobial agents include microorganisms, plants and synthesis^2^. Among the microbial sources, the leading antibiotics-producing bacteria are the actinomycetes which are a group of branching unicellular microorganisms and its dominating species is the *Streptomycetes* ^3, 4, 5^. Of the natural sources of antibiotics about 70% are isolated from actinomycetes e.g. Actinomycin (from *Actinomyces antibioticus*), Streptomycin (from *Streptomyces griseus*), Tetracycline (from *Streptomyces aureofaciens*), Chloramphenicol (from *Streptomyces venezuelae*), Vancomycin (from *Streptomyces orientalis*) and Gentamycin (from *Micromonospora purpurea* and *M. echinospora*) ^2, 6, 7^. The remaining 30% are from other microorganisms such as *Pseudomonas, Bacillus species* (bacitracin from *Bacillus subtilis* var Tracy; surfacins from *Bacillus subtilis* growing on molasses; inturinics, pumulin, bacilysin and gramicidin from *Bacillus subtilis*) and filamentous fungi e.g. penicillins and cephalosporins ^2, 6, 7^. Even though about 1,200 antibiotics have been isolated, only few have been commercially used for management and treatment of infections because of their toxicity ^2, 8^.

During the last few decades, the rate of production of novel antimicrobial agents from the source of terrestrial microorganism has declined until recently by the fact that extracts from soil-derived actinomycetes have yielded high numbers of clinically unacceptable metabolites ^9^. The aquatic environment is now witnessing a rich and unused pool of useful novel natural products. The marine environment alone is known to contain taxonomically diverse bacterial groups which exhibit unique physiological and structural characteristics that enable them to survive in extreme environmental conditions, with the potential production of novel secondary metabolites not observed in terrestrial microorganisms ^10^.

The environment in Ghana is full of an array of microorganisms whose antibiotic-producing potentials have not been adequately exploited. This study sought to investigate the possibility of identifying potent natural bioactive substances from this environment for antimicrobial drug development.

## Methods

### Sampling and Isolation of antimicrobial metabolites producing microorganisms

The Naa Korley: Sea and Lagoon at Korley Gonor in Greater Accra Region and two water bodies in Eastern Region of Ghana: River Densu and Nsukwao at Koforidua were selected for the study. Fourteen samples of water and soils were collected from each of these sites and conveyed to the laboratory on an ice. The water samples were collected by submerging sterile plastic containers into the water to a depth of about 10 cm and opened. They were filled with water and then closed before bringing to the surface. About five grams (5 g) each of soil samples were collected into sterile plastic containers with different sterilized spoons and labelled.

All samples were processed within 12 hours of collection. About 1 g of soil samples were suspended in 10 mL of sterile distilled water and shook. One milliliter (1 mL) of these suspensions were added to the molten nutrient agars (Oxoid). About one milliliter (1 mL) of water samples were also separately inoculated into a molten agar, mixed well, poured into sterile petri dish and allowed to solidify. The plates were incubated at 37 °C for seven days with daily observation. Eight (8) colonies were observed on each of the 14 agar plates incubated with a clear zone around them were carefully isolated into pure cultures.

### Test microorganisms

The test microorganisms used were obtained from the Department of Pharmaceutics Microbiology Laboratory KNUST, Kumasi and these included; *Staphylococcus aureus* (ATCC25923), *Streptococcus pyogenes* (clinical isolates), *Bacillus subtilis* (NCTC10073), *Pseudomonas aeruginosa* (ATCC27853), *Neisseria gonorrhoeae* (clinical isolate), *Escherichia coli* (NCTC9002), *Salmonella typhi* (NCTC 6017) and *Klebsiella pneumonia*e (Clinical isolate). The purpose for selecting the above bacteria was because they are pathogens commonly associated with bacterial infections.

### Screening of microbial isolates for antimicrobial metabolite production

The isolates were tested for their antimicrobial activity using the agar well diffusion method. One hundred and fifty-two (152) screwed capped test tubes containing 20 mL each of nutrient agar (Oxoid) were melted, stabilized at 45 °C for 15 minutes in water bath. Aseptically each of 0.1 mL of an 18-hour broth culture of test organisms (The density of these suspensions was adjusted to 0.5 McFarland standards) were inoculated into 19 different screw capped test tubes and transferred into sterile petri dishes under a laminar air flow cabinet, (Model T2 2472 Skan AG, Switzerland). After the agars plates were allowed to set, six wells were created equidistant from each other in the agar with cork borer (8 mm in diameter) and filled with 0.1 mL of 72 hours broth culture of an isolate microorganism. The plates were allowed to stand for one hour to allow diffusion of any metabolite in the broth into the seeded agar. They were then incubated at 37 °C for 24 hours and observed for signs of growth and zones of inhibition. The experiment was repeated two times and the diameters of the zones of inhibition were measured and recorded (Table 1). The isolates NKSEW_3_ and NKLS_6_ that showed high number of mean zone of growth inhibitions were selected for further studies.

**Table 1.** Antibiotic production screening of isolates from water and soil samples of River Densu

| Isolates | Test Microorganisms/ Average Zone of growth inhibition (mm) |  |  |  |  |  |  |  |
| --- | --- | --- | --- | --- | --- | --- | --- | --- |
|  | *S. aur* | *S. pyo* | *E. coli* | *P. aer* | *Gono* | *K. pne* | *B. Sub* | *S. typ* |
| ADW 1 | - | - | - | - | - | - | $29.5 \pm 0.5$ | $19.5 \pm 0.5$ |
| 2 | - | - | - | - | - | - | - | $21.5 \pm 0.5$ |
| 3 | - | - | - | - | - | - | $21.0 \pm 0.5$ | - |
| 4 | - | - | - | - | - | - | - | - |
| 5 | - | - | - | - | - | - | $20.5 \pm 0.5$ | - |
| 6 | - | - | - | - | - | - | $18.5 \pm 0.5$ | $19.5 \pm 0.5$ |
| 7 | - | - | - | - | - | - | $20.5 \pm 0.5$ | - |
| 8 | - | - | - | - | - | - | $18.5 \pm 0.5$ | $19.5 \pm 0.5$ |
| ODW 1 | - | - | - | - | $17.5 \pm 0.5$ | - | - | $20.5 \pm 0.5$ |
| 2 | - | - | - | - | $15.5 \pm 0.5$ | - | $20.5 \pm 0.5$ | - |
| 3 | - | - | - | - | $20.5 \pm 0.5$ | - | - | - |
| 4 | - | - | - | - | - | - | - | $15.5 \pm 0.5$ |
| 5 | - | - | - | - | 16.5± 0.5 | - | - | 15.5± 0.5 |
| 6 | - | - | - | - | 26.0± 0.5 | - | 20.5± 0.5 | 20.5± 0.5 |
| 7 | - | - | - | - | 16.5± 0.5 | - | 21.5± 0.5 | - |
| 8 | - | - | - | - | 20.5± 0.5 | - | 21.5± 0.5 | 20.5± 0.5 |
| WWDW<sub>1</sub>1 | - | - | - | 14.5± 0.5 | - | 14.5±0.5 | 19.5± 0.5 | 16.5±0.5 |
| 2 | - | - | - | 19.5± 0.5 | - | 14.5±0.5 | - |  |
| 3 | - | - | - | 16± 1.0 | - | 14.5±0.5 | - | 17.5±0.5 |
| 4 | - | - | - | 16.5± 0.5 | - | 18.5±0.5 | - | - |
| 5 | - | - | - | 18.5± 0.5 | - | 21.5±0.5 | 14.5± 0.5 | 17.5±0.5 |
| 6 | - | - | - | 20± 0.0 | 14.5±0.5 | 20.5±0.5 | - | 19.5±0.5 |
| 7 | - | - | - | 18.5± 0.5 | - | - | 14.5± 0.5 | - |
| 8 | - | - | - | - | 14.5±0.5 | - | - | 19.5±0.5 |
| WWDW<sub>2</sub>1 | - | - | - | - | 15.5± 0.5 | - | - | 26.5± 0.5 |
| 2 | - | - | - | - | 26.5± 0.5 | - | - |  |
| 3 | - | - | - | - | - | - | 19.5± 0.5 | 20.5± 0.5 |
| 4 | - | - | - | - | - | - | - | - |
| 5 | - | - | - | - | - | - | - | - |
| 6 | - | - | - | - | - | - | 14.5± 0.5 | - |
| 7 | - | - | - | - | - | - | 19.5±0.5 | - |
| 8 | - | - | - | - | - | - | - | - |
| WWDS<sub>1</sub>1 | - | - | - | 20.5±0.5 | - | - | 17.5 ± 0.5 | 24.5± 0.5 |
| 2 | - | - | 18.5±0.5 | - | - | - | 22.5 ± 0.5 | 16.5± 0.5 |
| 3 | - | - | - | - | - | - | 20.5 ± 0.5 | - |
| 4 | - | - | - | - | - | - | - | 21.5± 0.5 |
| 5 | - | - | - | 21.5±0.5 | - | - | 18.5 ± 0.5 | 21± 1 |
| 6 | - | - | 24.5±0.5 | - | - | - | 19.5 ± 0.5 | - |
| 7 | - | - | 21.5±0.5 | - | - | - | - | 16.5± 0.5 |
| 8 | - | - | - | 1.5±0.5 | - | - | 20.5 ± 0.5 | - |
| WWDS<sub>2</sub>1 | 14.5± 0.5 | 17.5± 0.5 | 15± 0.5 | - | 21± 1 | - | - | - |
| 2 | 16.5± 0.5 | - | - | - | - | - | - | - |
| 3 | - | 19± 0.5 | - | - | 24.5± 0.5 | - | - | - |
| 4 | - | - | 14.5± 0.5 | - | 24.5± 0.5 | - | - | - |
| 5 | 14.5± 0.5 | - | 18.5± 0.5 | - | 24.5± 0.5 | - | - | - |
| 6 | - | - | 18.5± 0.5 | - | 39.5± 0.5 | - | - | - |
| 7 | - | - | - | - | 24.5± 0.5 | - | - | - |
| 8 | 16.5± 0.5 | - | - | - | - | - | - | - |
| ODS 1 | - | - | - | - | - | - | - | - |
| 2 | - | - | - | - | - | - | - | - |
| 3 | - | - | - | - | - | - | - | - |
| 4 | - | - | - | - | - | - | - | - |
| 5 | - | - | - | - | - | - | - | - |
| 6 | - | - | - | - | - | - | - | - |
| 7 | - | - | - | - | - | - | - | - |
| 8 | - | - | - | - | - | - | - | - |
| ADS 1 | - | - | 35.5± 0.5 | - | 19.5± 0.5 | 36± 1 | - | 38.5± 0.5 |
| 2 | - | - | 35.5± 0.5 | - | - | 36.5± 0.5 | - | 39.5± 0.5 |
| 3 | - | - | 35.5± 0.5 | - | - | 38.5± 0.5 | - | 38± 1 |
| 4 | - | - | 26.5± 0.5 | - | - | 38.5± 0.5 | - | 38.5± 0.5 |
| 5 | - | - | 26.5± 0.5 | - | - | 38.5± 0.5 | - | 35.5± 0.5 |
| 6 | - | - | 25.5± 0.5 | - | - | 36± 1 | - | 39.5± 0.5 |
| 7 | - | - | 26.5± 0.5 | - | - | - | - | - |
| 8 | - | - | - | - | - | 38.5± 0.5 | - | - |

ODW = River Densu Wobotumpan water, ADW = River Densu Akwadum water, WWDW<sub>1</sub> = River Densu Waterworks number one water, WWDW<sub>2</sub> = River Densu Waterworks number two water, WWDS<sub>1</sub> = River Densu Waterworks number one soil, WWDS<sub>2</sub> = River Densu Waterworks number two soil, ODS = River Densu Wobotumpan soil, ADS = River Densu Akwadum soil, *S. aur* = *Staphylococcus aureus* (ATCC25923), *S. pyo* = *Streptococcus pyogenes* (clinical isolates), *E. Col* = *Escherichia coli* (NCTC9002), *P. aer* = *Pseudomonas aeruginosa* (ATCC27853), *N. Gono* = *Neisseria gonorrhoeae* (clinical isolate), *K. pne* = *Klebsiella pneumoniae* (Clinical isolate), *B. Sub* = *Bacillus subtilis* (NCTC10073) and *S. typ* = *Salmonella typhi* (NCTC 6017)

### Effect of some growth factors on antimicrobial metabolite production

#### Time of Incubation

Two (2) milliliters of the isolates NKSEW_3_ and NKLS_6_ were selected to produce high zones of growth inhibitions against at least three of the test organisms were inoculated into two test tubes containing 20 mL of sterile nutrient broth and incubated for 12 days at 37 °C. Samples were collected daily and tested against *Pseudomonas aeruginosa* (ATCC27853), *Escherichia coli* (NCTC9002) and *Klebsiella pneumonia*e (clinical isolate) for antimicrobial activity using agar well diffusion method.

#### Temperature

About 1 mL of the broth cultures of isolates NKSEW_3_ and NKLS_6_ were separately inoculated into 10 mL nutrient broths and incubated at different temperatures of 20 °C, 25 °C, 34 °C, 37 °C and 45 °C for 72 hours. They were then centrifuged at 3622 g for 1 h to precipitate the microbial cells from the metabolite solutions. The metabolite solutions obtained were then analyzed for antimicrobial activity against *Pseudomonas aeruginosa* (ATCC27853), *Escherichia coli* (NCTC9002) and *Klebsiella pneumonia*e (clinical isolate).

#### Carbon and Nitrogen sources

The effect of carbon sources on antimicrobial metabolite production was evaluated by growing the isolate in 10 mL fermentation media fortified with 60 mg of various carbon compounds: glucose, galactose, xylose, starch and glycerol). The metabolite solutions obtained were tested for antimicrobial activity against *Pseudomonas aeruginosa* (ATCC27853), *Escherichia coli* (NCTC9002) and *Klebsiella pneumonia*e (clinical isolate). The procedure was repeated for nitrogen compounds: sodium nitrate, potassium nitrate, ammonium chloride, ammonium nitrate and ammonium sulphate) ^11^.

#### pH

The optimum pH for maximum antimicrobial activity was evaluated by fermenting isolate NKSEW_3_ and NKLS_6_ in tubes of 10 mL nutrient broth at varying pHs (4, 6, 7, 8 and 9) after which their antibacterial activity was evaluated by the cup plate method as above ^11^.

### Fermentation and extraction of metabolites of NKSEW_3_ and NKLS_6_

The isolate NKSEW_3_ was inoculated into 2 L of nutrient broth and incubated at 37 °C for 10 days. The culture was then centrifuged at 3622 g for 1 hour and the supernatant filtered, extracted with ethyl acetate. It was transferred into a separation funnels and allowed to stand for 1 hour in order to facilitate good separations. The two distinct formed layers (aqueous at the bottom and the organic at the top of the separating funnel) were evaluated for antimicrobial activity using disk diffusion method. The aqueous layer showed considerable growth inhibition and was evaporated to dryness and dried at room temperature (25°C). Two replicates were done and the extracts obtained were weighed and kept in a desiccator for use. The procedure was repeated for isolate NKLS_6_ ^11^.

### Determination of MIC and MBC of the extracts

Minimum Inhibitory Concentration (MIC) was determined using the broth dilution method ^11^. Serial dilutions (80 μL) of the extract in nutrient broth (Sigma-Aldrich Chemie, GmbH) in the range of 0.78 mg/mL to 100 μg/mL were made in 96-well micro-plates (Thermo Scientific, USA). The inocula (20 μL) of the test microorganisms prepared from 18 h broth cultures (containing 10^5^ cfu/mL) were dispensed into the plates. Three replicates were made. The plates were incubated at 37 °C for 24 hours. Bacterial growth was determined after addition of 20 μL of 0.2 mg/mL MTT (Sigma-Aldrich, St. Louis, MO, USA). The minimum bactericidal concentration (MBC) test was performed as above in the MIC determination except that 80 μL aliquots were withdrawn from wells that showed inhibition in the MIC experiment and inoculated into 5 mL nutrient broths. These were incubated at 37 °C for 48 hours and observed for signs of growth.

### Fractionation of the extracts of isolates NKSEW_3_ and NKLS_6_

Column chromatography was employed to fractionate the extracts. Silica gel G (mesh size 4-30 μm) was used as stationary phase for the column chromatography. Forty grams (40 g) of silica was weighed into a beaker and 150 mL of petroleum ether was added to form a slurry. This was transferred into a glass column that had been washed and rinsed with petroleum ether. More of the petroleum ether was added to the column and allowed to drain through the settled silica until it became the same in appearance indicating a well packed column.

Ten grams (10 g) of the extract of isolate NKSEW_3_ was dissolved in 300 mL of methanol. It was then absorbed on 10 g of silica and the solvent was evaporated. The sample was applied on the top bed and covered with absorbent cotton. The column was then run with a gradient elution system of petroleum ether, ethyl acetate and methanol. A total of 10 aliquots made up of 100 mL each was collected and evaporated at room temperature to dryness and weighed. The fractions were subjected to TLC and only three aliquots contained a compound. Aliquots with similar TLC profiles were bulked together to form fraction NKSEW_3_1.

Similarly, ten grams (10 g) of the extract of isolate NKLS_6_ was dissolved in 300 mL of methanol. It was then absorbed on 10 g of silica and the solvent was evaporated. The sample was applied on the top bed and covered with absorbent cotton. The column was run with a gradient elution system of petroleum ether, ethyl acetate and methanol. A total of 10 aliquots made up of 100 mL each was collected and evaporated at room temperature to dryness and weighed. The fractions were subjected to TLC and only four aliquots contained a compound. Aliquots of similar TLC profiles were bulked together to form fraction NKLS_6_2.

### Antimicrobial activity of the fractions NKSEW_3_1 and NKLS_6_2

The antimicrobial activity of the fractions NKSEW_3_1 and NKLS_6_2 were determined using disk diffusion method. Sterile filter disks (6 mm) were saturated separately with the fractions. They were air dried and placed on the seeded agar plates (1×10^8^ CFU/mL) of *Pseudomonas aeruginosa* (ATCC 27853), *Escherichia coli* (NCTC9002) and *Klebsiella pneumonia*e (clinical isolate). They were incubated at 37 °C for 24 hours. The plates were observed for zone of growth inhibitions and they were measured in millimeters.

### High Performance Liquid chromatography (HPLC) for the fraction NKLS_6_2

The fraction NKSL_6_2 was subjected to High performance liquid chromatography (HPLC) using uBoundapat C18 column (3.9 × 300 mm, 10 μm particle size, 125 Ǻ with maximum pressure of 2000 psi) and a system software chromeleon. Gynkotek pump embedded with a degasser of Dionex DG-1210, a detector of Dionex UVD 340S and a sampler of Dionex Gina 50 were the set up used while the separation of the mixture, the percentage of methanol-water solvent system was used.

### Gas Chromatography–Mass Spectrometry (GC-MS) analysis of the fraction NKLS_3_2

Identification of the chemical compounds present in the fraction of the extract was executed by GC-MS. Analysis was performed on a factor four^TM^ capillary column (30.0 m × 250 μm; Varian, Middelburg, The Netherlands) under the following conditions: carrier gas of Helium, 2.00 min as solvent delay, 150 °C and 250 °C as source and transfer temperatures respectively, 0.1 µL as volume, and 250 °C of injection, and 20:1 as split ratio initially and closed at 0.00 min, and open again (30:1) at 26 minutes. The initial temperature of the oven was set at 40 °C at 0 min and then 3 °C/min ramp to 90 °C which was held at 0 minute; again 10 °C/min ramp to 240 °C. The mass spectrometer transfer line temperature was 280 °C with ion trap and manifold temperatures as 220 °C and 120 °C respectively. The full scan was 40-450 Da) EI auto mode with filaments of 20 µA was employed for the MS analysis from 5.00-28.00 min which gave 0.78 s/scans of 3 µ scan. The automatic target control and the multiplier voltage were 20,000 and 1450 V respectively. The baseline offset, the peak found with S/N of the quantifier ion, and the peak width were 5, 3 and 2 s respectively was set as the parameters for the processing of the peaks in the chromatograms. The National Institute of Standard Technology (NIST) library spectra to match with minimum similarity, it was kept at 500 reversed fit. The basis of diagnostic ion, the peak assignments and integration were the quantification and was determined automatically using thermomass software

## Results

### Isolation of antibiotic producing organisms

A total of 112 isolates of microorganisms were obtained out of fourteen samples (River Densu (8), River Nsukwao (2) at Koforidua, Nii Korley Sea (2) and Nii Korley Lagoon (2) at Accra). Eight isolates were picked from each of the sample cultivated on the nutrient agar based on the clear zones around their colonies (Tables 1 - 3). The agar well diffusion method was used to screen the isolates for antibiotic production. The mean zone of growth inhibitions obtained were due to the antimicrobial agents’ presence in the isolates. The mean zones of growth inhibition of 26 isolates ranged between 14.5±0.5 - 39.5± 0.5 mm against at least three of the test organisms (Tables 1 - 3).

**Table 2.** Antibiotic production screening of isolates from River Nsukwao and soil samples

| Isolates | Test Microorganisms/ Average Zone of growth inhibition (mm) |  |  |  |  |  |  |  |
| --- | --- | --- | --- | --- | --- | --- | --- | --- |
|  | *S. aur* | *S. pyo* | *E. coli* | *P. aer* | *Gono* | *K. pne* | *B. Sub* | *S. typ* |
| NKFW 1 | - | - | - | 24± 0.5 | - | - | - | 26.5± 0.5 |
| 2 | - | - | - | 23± 0.5 | - | - | - | - |
| 3 | - | - | - | - | 26.5± 0.5 | - | - | 19.5± 0.5 |
| 4 | - | - | - | 29.5± 0.5 | - | - | - | - |
| 5 | - | - | - | - | - | - | - | - |
| 6 | - | - | - | - | 21.5± 0.5 | - | - | 16.5± 0.5 |
| 7 | - | - | - | - | 21.5± 0.5 | - | - | 16.5± 0.5 |
| 8 | - | - | - | 29.5±0.5 | - | - | - | - |
| NKFS 1 | 16.5± 0.5 | - | - | 18.5± 0.5 | - | - | - | - |
| 2 | - | - | - | - | - | - | 14.5± 0.5 | - |
| 3 | - | - | - | - | - | - | - | - |
| 4 | - | - | - | - | - | - | - | - |
| 5 | 14.5± 0.5 | - | 14.5± 0.5 | - | - | - | - | - |
| 6 | - | - | - | - | - | - | - | - |
| 7 | - | 14.5± 0.5 | - | - | - | - | - | - |
| 8 | - | - | - | 14.5± 0.5 | - | - | - | - |

NKFW = Nsukwao Koforidua water, NKFS = Nsukwao Koforidua soil, *S. aur* = *Staphylococcus aureus* (ATCC25923), *S. pyo*= *Streptococcus pyogenes* (clinical isolates), *E. Col*= *Escherichia coli* (NCTC9002), *P. aer*= *Pseudomonas aeruginosa* (ATCC27853), *N. Gono*= *Neisseria gonorrhoeae* (clinical isolate), *K. pne*= *Klebsiella pneumoniae* (Clinical isolate), *B. Sub*= *Bacillus subtilis* (NCTC10073) and *S. typ*= *Salmonella typhi* (NCTC 6017).

**Table 3.** Antibiotic production screening of isolates from Nii Korley water and soil samples

| Isolates | Test Microorganisms/ Average Zone of growth inhibition (mm) |  |  |  |  |  |  |  |
| --- | --- | --- | --- | --- | --- | --- | --- | --- |
|  | *S. aur* | *S. pyo* | *E. coli* | *P. aer* | *Gono* | *K. pne* | *B. Sub* | *S. typ* |
| NKLS 1 | 18.5± 0.5 | - | - | - | 20.5± 0.5 | - | - | 26.5± 0.5 |
| 2 | 19.5± 0.5 | - | - | - | - | - | 15.5± 0.5 | - |
| 3 | - | - | - | - | 25.5± 0.5 | - | 15.5± 0.5 | 24.5± 0 |
| 4 | - | - | - | - | 18.5± 0.5 | - | - | - |
| 5 | - | - | - | - | 15.5± 0.5 | - | - | - |
| 6 | - | - | 19.5± 0.5 | 25.5± 0.5 | - | 27.5±0.5 | - | - |
| 7 | 16.5± 0.5 | - | - | - | - | - | - | - |
| 8 | - | - | - | 16.5± 0.5 | - | - | - | - |
| NKLW 1 | - | - | - | 18.5± 0.5 | - | 20.5±0.5 | - | 16.5± 0.5 |
| 2 | - | - | - | 18.5± 0.5 | - | - | - | - |
| 3 | - | - | - | - | 36.5± 0.5 | - | - | - |
| 4 | - | - | - | 29.5± 0.5 | - | - | - | - |
| 5 | - | - | - | 16.5± 0.5 | - | 24.5±0.5 | - | - |
| 6 | - | - | - | - | 18.5± 0.5 | - | - | - |
| 7 | - | - | 18.5± 0.5 | - | - | 24.5±0.5 | - | - |
| 8 | 18.5± 0.5 | - | - | 24.5±0.5 | - | - | - | - |
| NKSES 1 | - | - | - | - | - | 18.5± 0.5 | - | - |
| 2 | - | 16.5± 0.5 | - | - | - | - | - | - |
| 3 | - | 18.5± 0.5 | 14.5± 0.5 | - | - | - | - | - |
| 4 | - | - | - | - | - | - | 15.5± 0.5 | - |
| 5 | - | - | 19.5± 0.5 | - | - | - | - | 15.5± 0.5 |
| 6 | - | - | 18.5± 0.5 | - | - | - | 16.5± 0.5 | 17.5± 0.5 |
| 7 | - | - | - | - | 18.5± 0.5 | - | - | - |
| 8 | 14.5± 0.5 | - | 18.5± 0.5 | - | - | - | - | - |
| NKSEW1 | - | 18.5± 0.5 | - | - | 14.5± 0.5 | - | 16.5± 0.5 | - |
| 2 | - | - | - | - | - | - | - | - |
| 3 | - | - | - | - | 16.5± 0.5 | - | 20.5± 0.5 | 18.5± 0.5 |
| 4 | 15.5± 0.5 | - | - | - | - | - | 19.5± 0.5 | 14.5± 0.5 |
| 5 | - | 15.5± 0.5 | - | - | - | - | - | - |
| 6 | 14.5± 0.5 | - | - | - | - | - | - | - |
| 7 | 14.5± 0.5 | - | - | - | - | - | 14.5± 0.5 | - |
| 8 | - | - | 14.5± 0.5 | - | - | - | - | 16.5± 0.5 |

NKLS = Nii Korley Lagoon soil, NKLW = Nii Korley Lagoon water NKSES = Nii Korley sea soil, NKSEW = Nii Korley sea water, *S. aur*= *Staphylococcus aureus* (ATCC25923), *S. pyo*= *Streptococcus pyogenes* (clinical isolates), *E. Col*= *Escherichia coli* (NCTC9002), *P. aer*= *Pseudomonas aeruginosa* (ATCC27853), *N. Gono*= *Neisseria gonorrhoeae*(clinical isolate), *K. pne*= *Klebsiella pneumoniae* (Clinical isolate), *B. Sub*= *Bacillus subtilis* (NCTC10073) and *S. typ*= *Salmonella typhi* (NCTC 6017).

It was recorded that 40 isolates showed zone of growth inhibition against *S. typhi,* 31 against *Gonococcus*, 24 against *P. aeruginosa*, 19 against *K. pneumonia*, 17 against *S. aureus,* 16 against *B. subtilis,* 15 against *E. coli* and 7 against *S. pyogenes* while the isolates ODS (1-8) showed no zone of growth inhibition when their metabolites were screened against the test organisms. The isolates NKSEW_3_ and NKLS_6_ were chosen for secondary screening since they showed relatively high zones of growth inhibitions (18.5± 0.5 - 26.5±0.5 mm for NKSEW_3_ and 19.5±0.5 - 27.5±0.5 mm for NKLS_6_ (Table 1 - 3) and (Plate 1)

**Plate 1.**
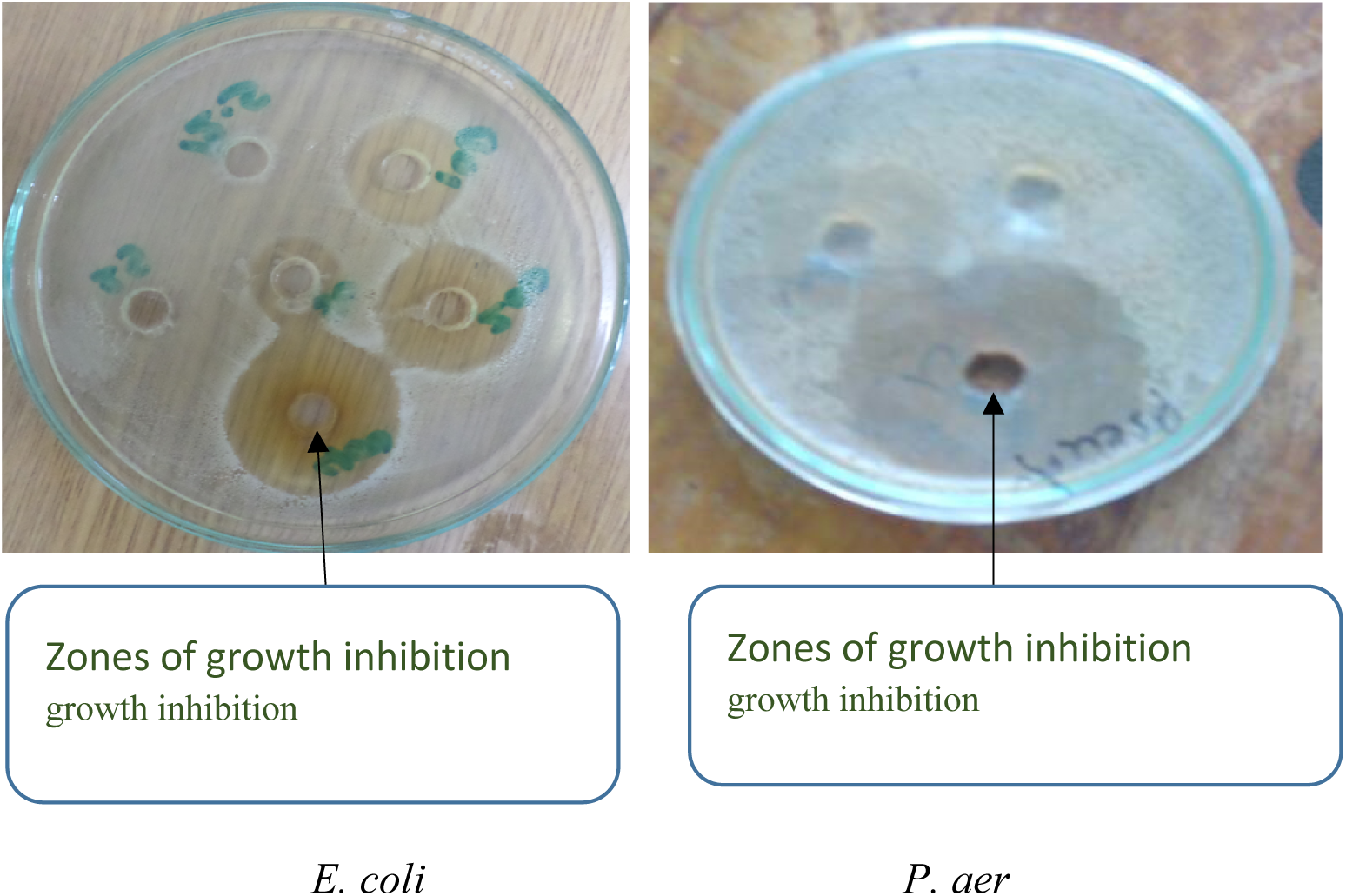
Some of the isolates zones of growth inhibition produced during screening.

### Incubation period for optimum Antimicrobial activity

The selected isolates NKSEW_3_ and NKLS_6,_ when incubated and tested daily for 12 days showed growth inhibition of *E. coli, P. aeruginosa* and *K. pneumoniae*. The highest zones of growth inhibition were obtained on days 9, 10 and 11. On day 12, the growth inhibition started declining (Figure 1).

**Figure 1.**
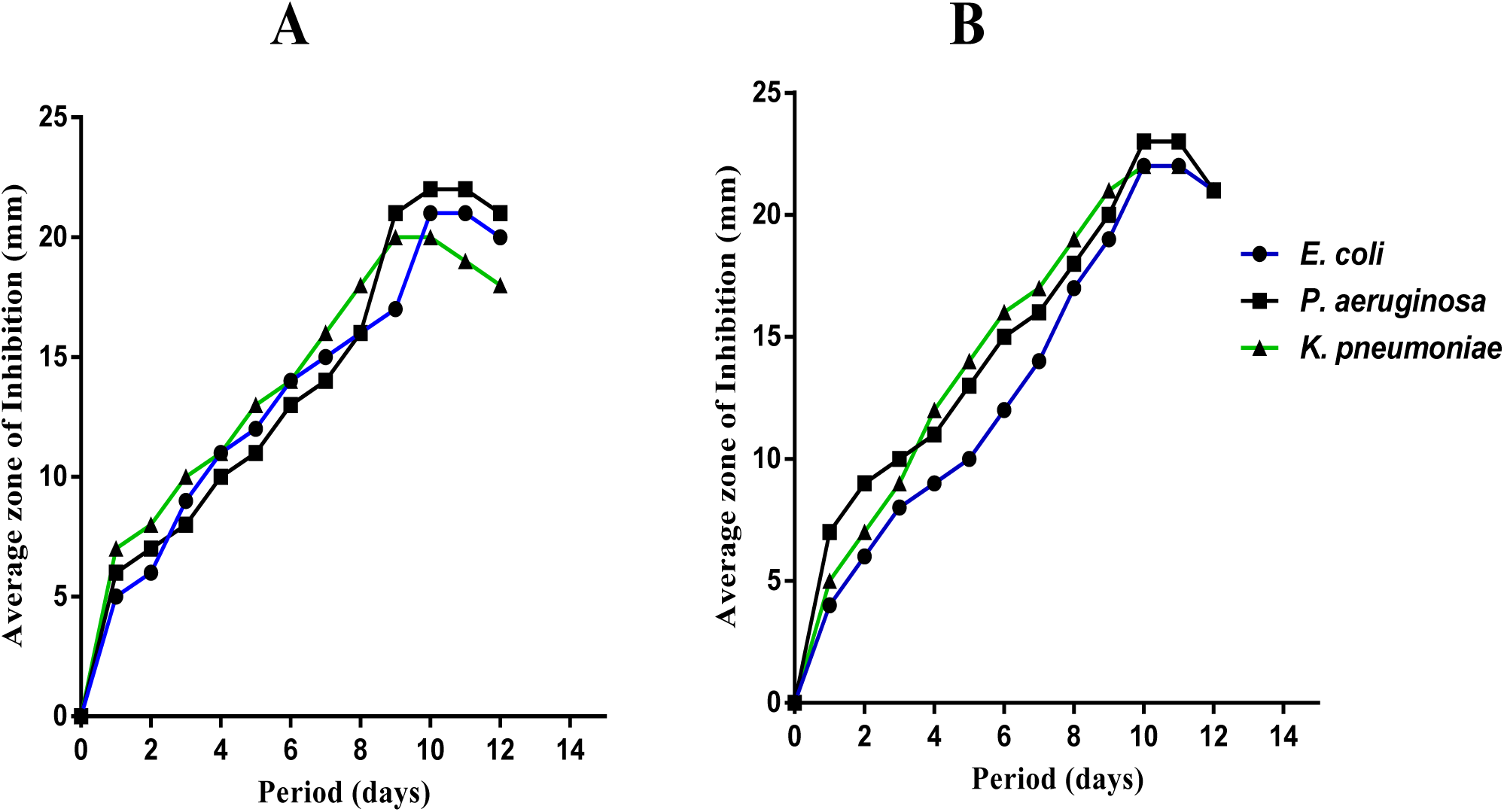
Effect of length of incubation on antibiotic producing organisms of isolates NKSEW3 (A) and NKLS6 (B).

**Figure 2.**
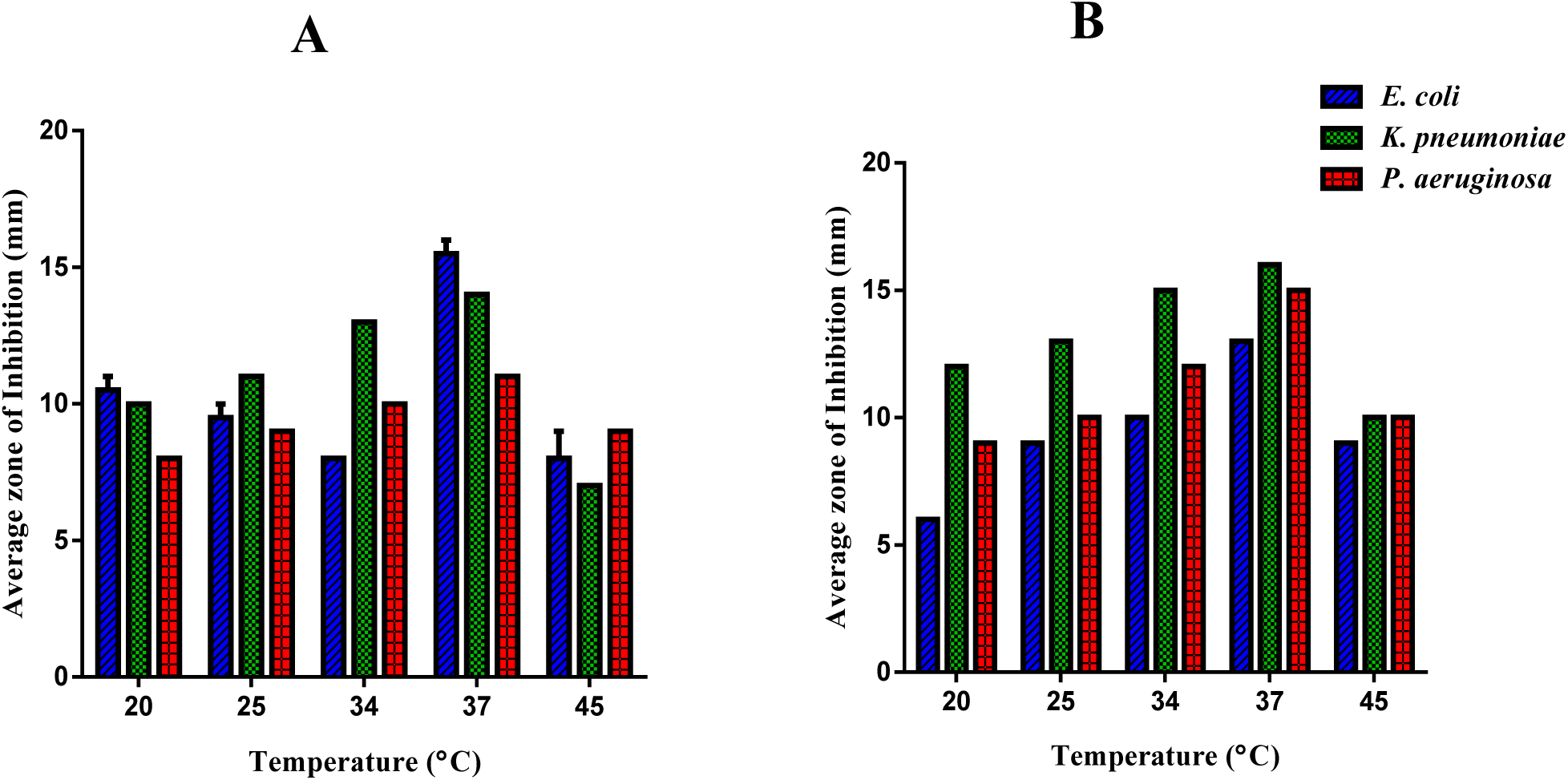
Effect of Temperature on antibiotic producing organisms of isolates (A) NKSEW_3_ and (B) NKLS_6_

**Figure 3.**
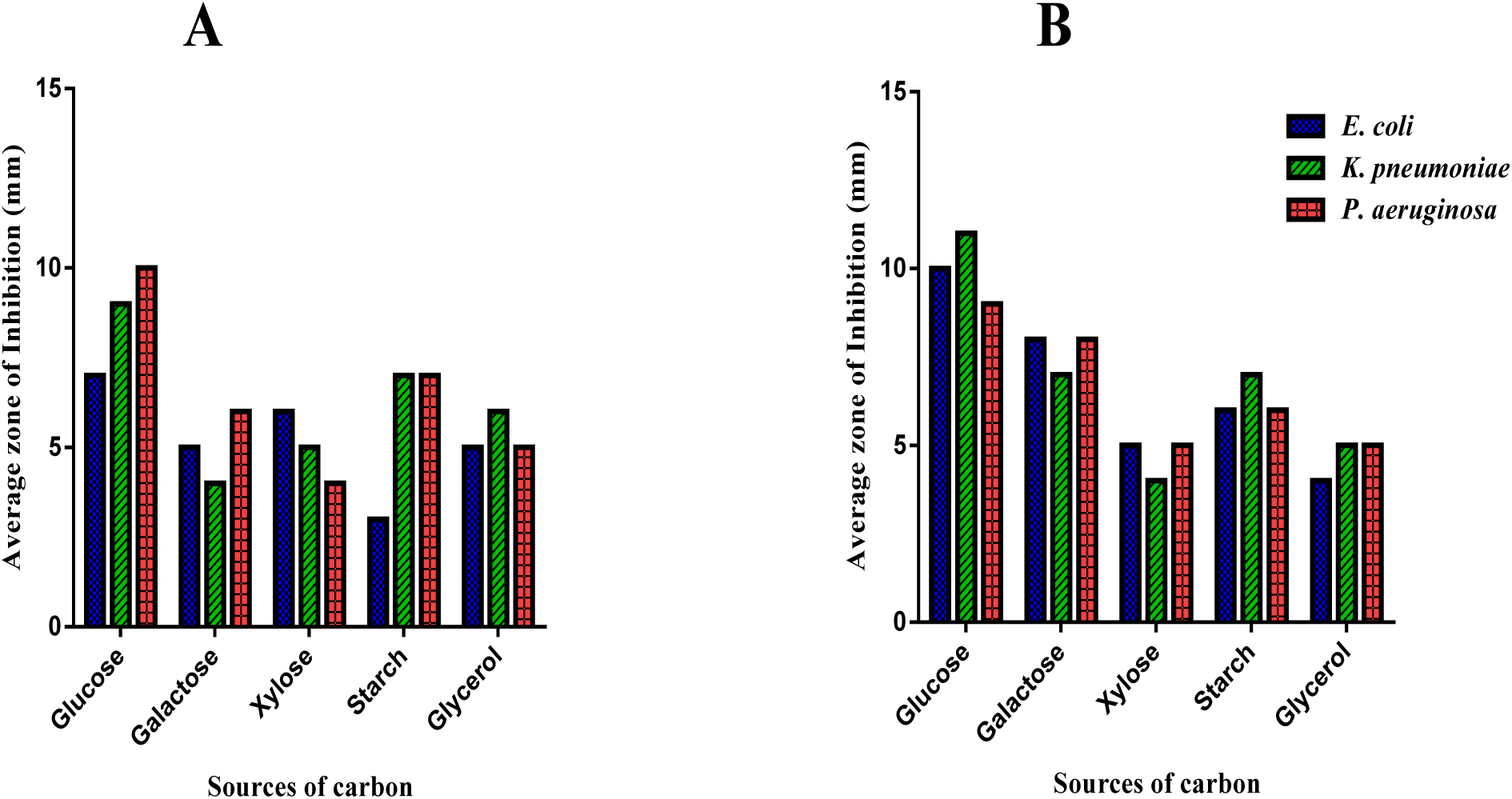
Effect of sources of Carbon on antibiotic producing organisms of isolates NKSEW_3_ (A) and NKLS_6_. (B)

**Figure 4.**
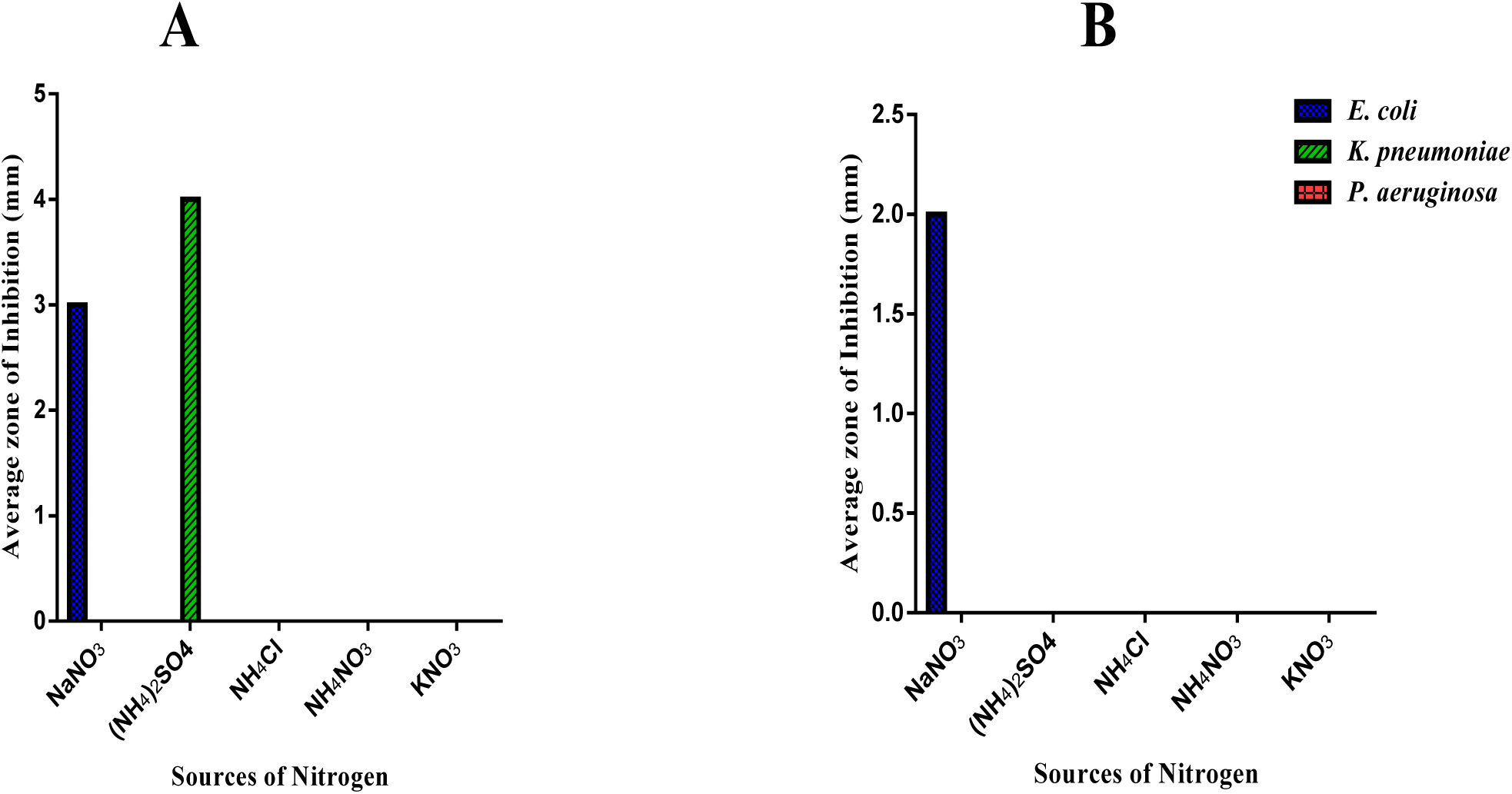
Effect of sources of Nitrogen on antibiotic producing organisms of isolates NKSEW_3_ (A) and NKLS_6_. (B).

### Extraction and Antimicrobial activity testing of the extracts of isolates NKLS_6_ and NKSEW_3_

The isolates NKLS_6_ and NKSEW_3_ were selected based on the results of their growth of inhibitions against the test organisms recorded (Table 3). The selected isolates NKLS_6_ and NKSEW_3_ were extracted using ethyl acetate and they produced hygroscopic brownish precipitate after evaporation. The antimicrobial activity testing of the extracts of isolates NKLS_6_ and NKSEW_3_ using agar well diffusion method showed zone of growth inhibition against. *E. coli, P. aeruginosa* and *K. pneumoniae*. The mean zones of growth inhibitions ranged between 18.00±0.50 - 20.50±0.50 mm and 12.00±0.00 - 15.00±0.50 mm at the concentration 100 mg/mL. At a concentration of 50 mg/mL against the test organisms ranged between 15.00±0.50 - 18.00±0.00 mm and 8.50±0.50 - 11.00±0.00 mm whiles at a concentration 25 mg/mL, the zones of growth inhibition were observed ranged between 7.50±0.50 - 10.50±0.50 mm and 6.00±0.50-7.50±0.50 mm for extracts NKLS_6_ and NKSEW_3_ respectively. It was recorded that the antimicrobial activity of the extracts are in dose-dependent manner. The minimum and maximum average zones of growth inhibitions of ciprofloxacin (1 mg/mL) against the test organisms ranged between 33.00±0.50 and 43.00±0.00 mm (Table 4).

**Table 4.**
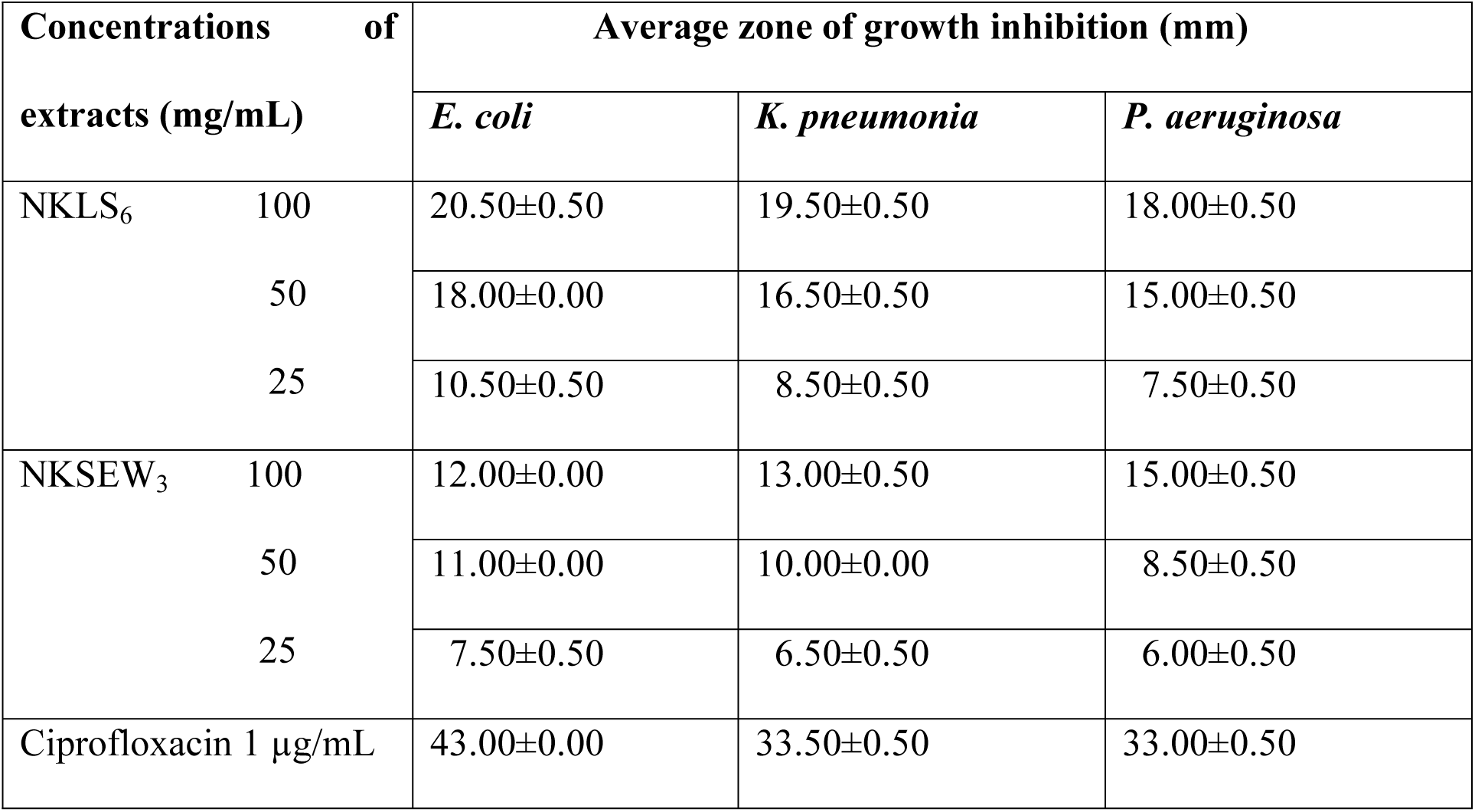
Antimicrobial activities of extracts of isolates NKSEW3; NKLS6 and Ciprofloxacin

### Minimum Inhibitory Concentrations (MIC) and Minimum Bactericidal Concentrations (MBC) of extracts of isolates NKSEW3, NKLS6 and Ciprofloxacin

Two test organisms had their MICs at 12.50 mg/mL for NKSEW_3_ and NKLS_6_; two had theirs at 6.25 mg/mL and one at 3.13 mg/mL for extracts NKSEW_3_ and NKLS_6_ (Table 5). The extracts were kill to *Escherichia coli and Pseudomonas aeruginosa* at concentration 25 mg/mL, whiles *K. pneumoniae* was killed at 12.50 mg/mL and 6.25 mg/mL for extracts of isolates NKSEW_3_ and NKLS_6_ respectively. With ciprofloxacin, *Escherichia coli and Klebsiella pneumoniae* had the highest MIC at 0.75 μg/mL followed by *Pseudomonas aeruginosa* at 0.38 μg/mL. Ciprofloxacin was able to *Escherichia coli* at 0.75 μg/mL while *Klebsiella pneumonia* and *Pseudomonas aeruginosa* were killed at 1.50 μg/mL (Table 5)

**Table 5.** MICS and MBCS of extracts and Ciprofloxacin

| Test Organisms | Extract NKSEW<sub>3</sub> (mg/mL) |  | Extract NKLS<sub>6</sub> (mg/mL) |  | Ciprofloxacin (µg/mL) |  |
| --- | --- | --- | --- | --- | --- | --- |
|  | MICs | MBCs | MICs | MBCs | MICs | MBCs |
| *E. coli* | 12.50 | 25.00 | 6.25 | 12.50 | 0.75 | 0.75 |
| *K. pne* | 6.25 | 12.50 | 3.13 | 6.25 | 0.75 | 1.5 |
| *P. aer* | 12.50 | 25.00 | 12.50 | 25.00 | 0.38 | 1.50 |

*E. coli*= *Escherichia coli*, *K. pne* = *Klebsiella pneumoniae* *P. aer* = *Pseudomonas aeruginosa*

### Effect of pH on isolates NKSEW3 and NKLS6 for antibiotic production

The isolates NKSEW_3_ and NKLS_6_ were grown at various initial pH values at 37 °C. There were lowest growth activity observed at pH values of 4, 6 and 9 whiles at pH value of 8 there were higher activities against *E. coli, P. aeruginosa* and *K. pneumoniae* for isolates NKSEW_3_ and NKLS_6_ (Figure 5).

**Figure 5.**
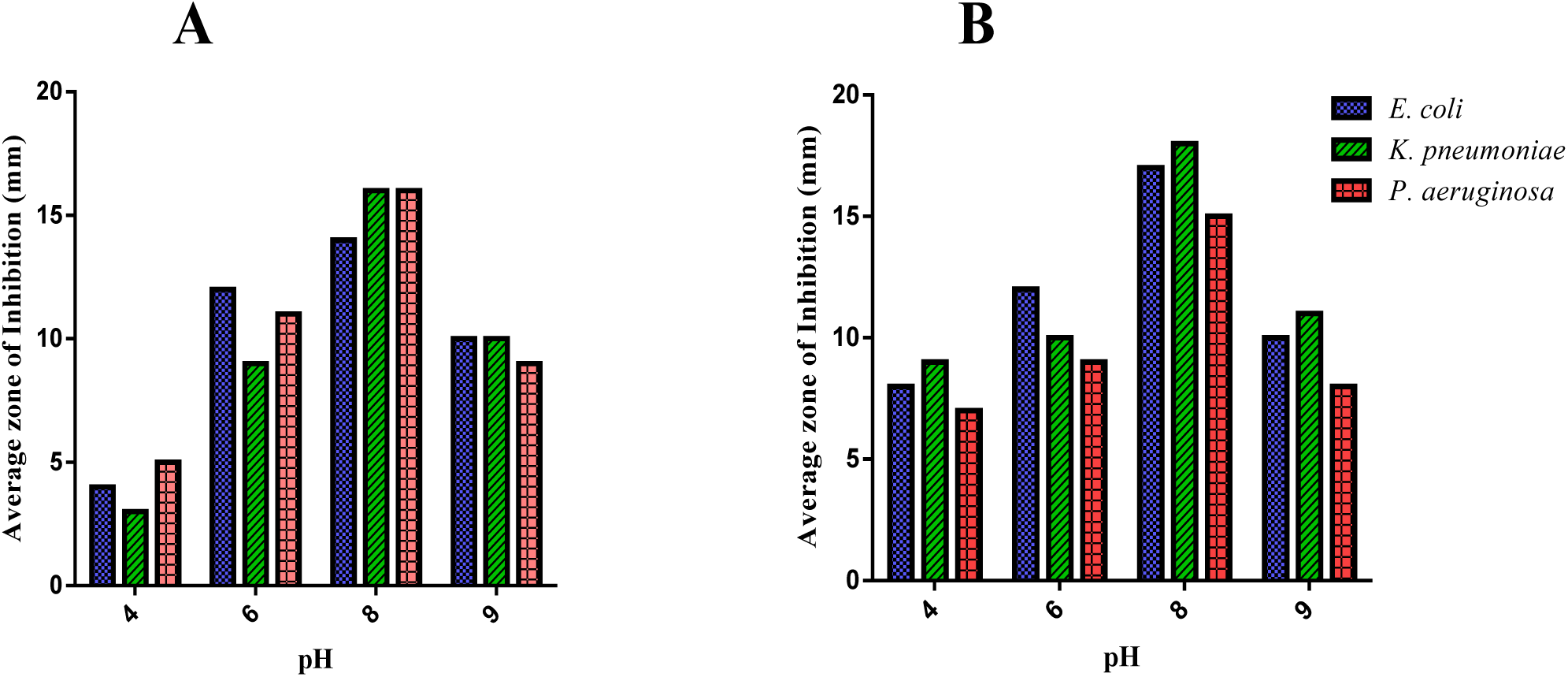
Effect of pH on antibiotic producing organisms of isolates NKSEW_3_ (A) and NKLS_6_ (B)

### Characterization of isolates NKSEW_3_ and NKLS_6_

The Gram staining of the isolates NKSEW_3_ and NKLS_6_ showed that the organisms were both Gram negative cocci and identified as *Pseudomonas species* using chromogenic medium (chromo agar). The biochemical tests of the isolates NKSEW_3_ and NKLS_6_ did not produce gases in the fermentation media but rather acids upon addition of glucose and lactose. It was also observed that both isolates grow at 42 °C and 6.5% NaCl but could not grow on cetrimide agar, macConkey agar, mannitol salt agar and bismuth sulphide agar as well as citrate, hydrogen sulphide, indole and nitrate.

### Fractionation of the Extracts of Isolates NKSEW_3_ and NKLS_6_

A total of 10 aliquots made up of 100 mL each of the extracts were collected and evaporated at room temperature to dryness. The fractions were subjected to TLC and only five aliquots contained a compound. Aliquots of similar TLC profiles were bulked together with retardation factors (R_f_) values as 0.56 for fraction NKSEW_3_1 and 0.44 for fraction NKLS_6_2. Antimicrobial activity of the fractions was evaluated using the disk diffusion method. Different sterile filter disks (6 mm) were saturated separately with the fractions in different sterile containers. They were air dried and placed on the seeded agar plates (1×10^8^ CFU/mL) of *Pseudomonas aeruginosa* (ATCC 27853), *Escherichia coli* (NCTC9002) and *Klebsiella pneumonia*e (clinical isolate). They were incubated at 37 °C for 24 hours. The plates were observed for zones of growth inhibitions and they were measured in millimeters (Table 6).

**Table 6.**
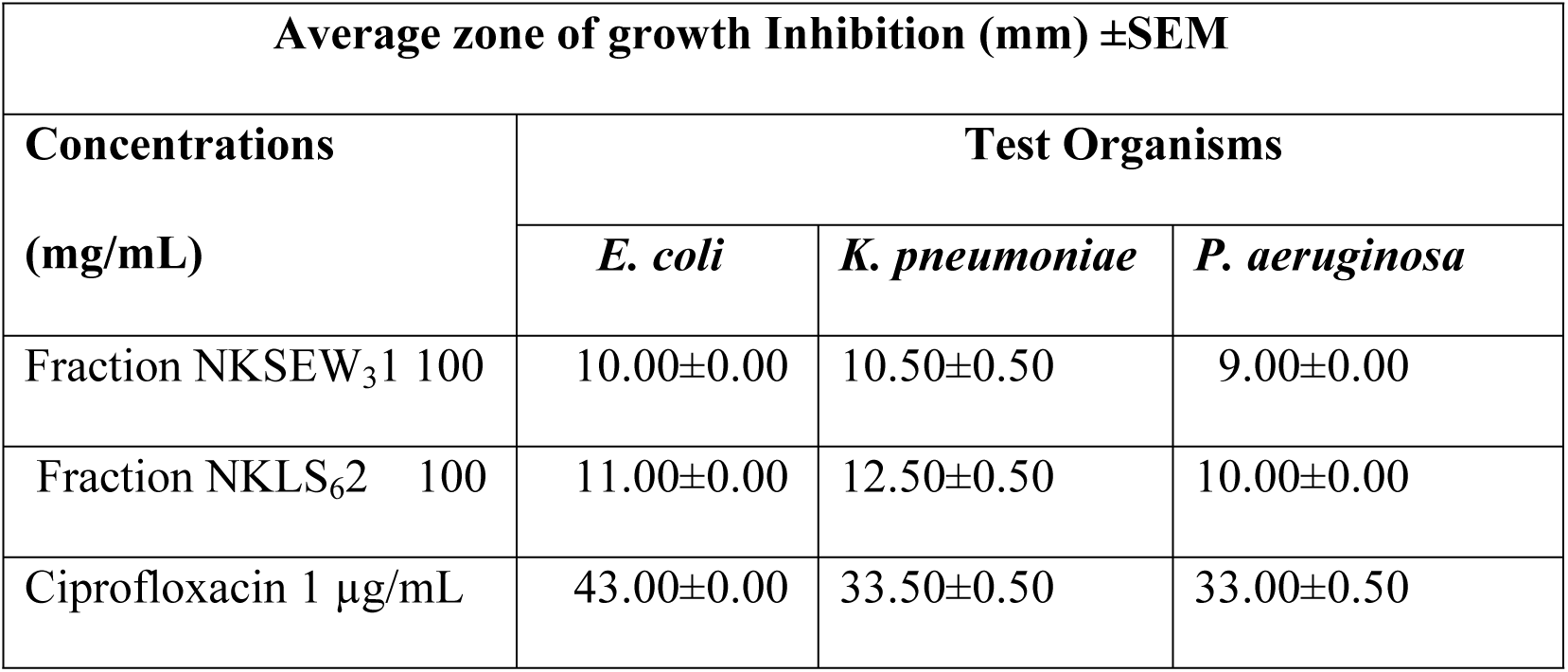
Test for the antimicrobial activity of the fractions NKSEW_3_1 and NKLS_6_2

### HPLC analysis of fraction NKLS_6_2

The HPLC profile of the fraction NKLS_6_2 revealed four peaks implying that the fraction is likely to contain four compounds (Figure 6).

**Figure 6.**
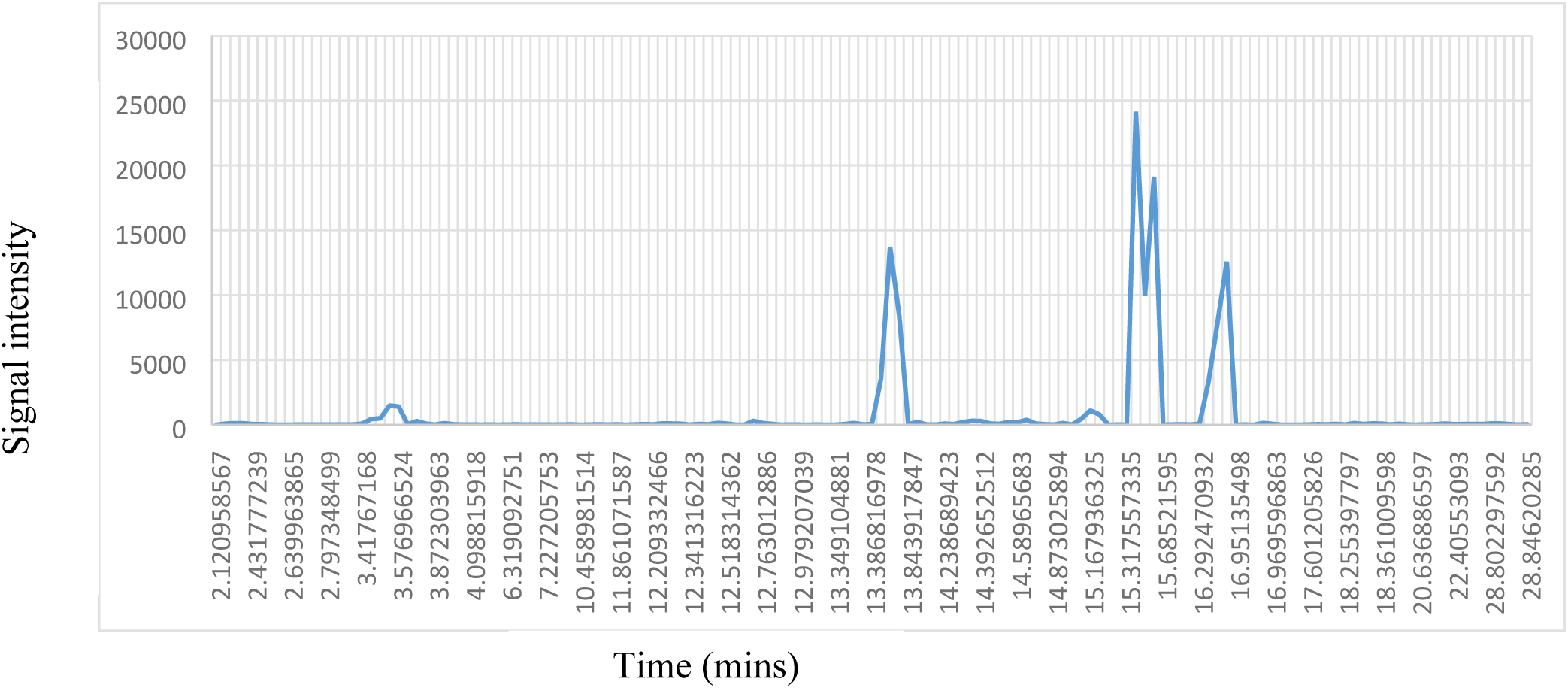
HPLC results of fraction NKLS_6_2

### Gas Chromatography –Mass Spectrometry (GC-MS) Analysis of the fraction NKLS_3_2

The GC-MS chromatogram of the NKLS_6_2 fraction revealed five compounds with their molecular formula, retention time (min.), molecular weight, peak area (%) and chemical structures (Table 7 and figure 7-12).

**Figure 7.**
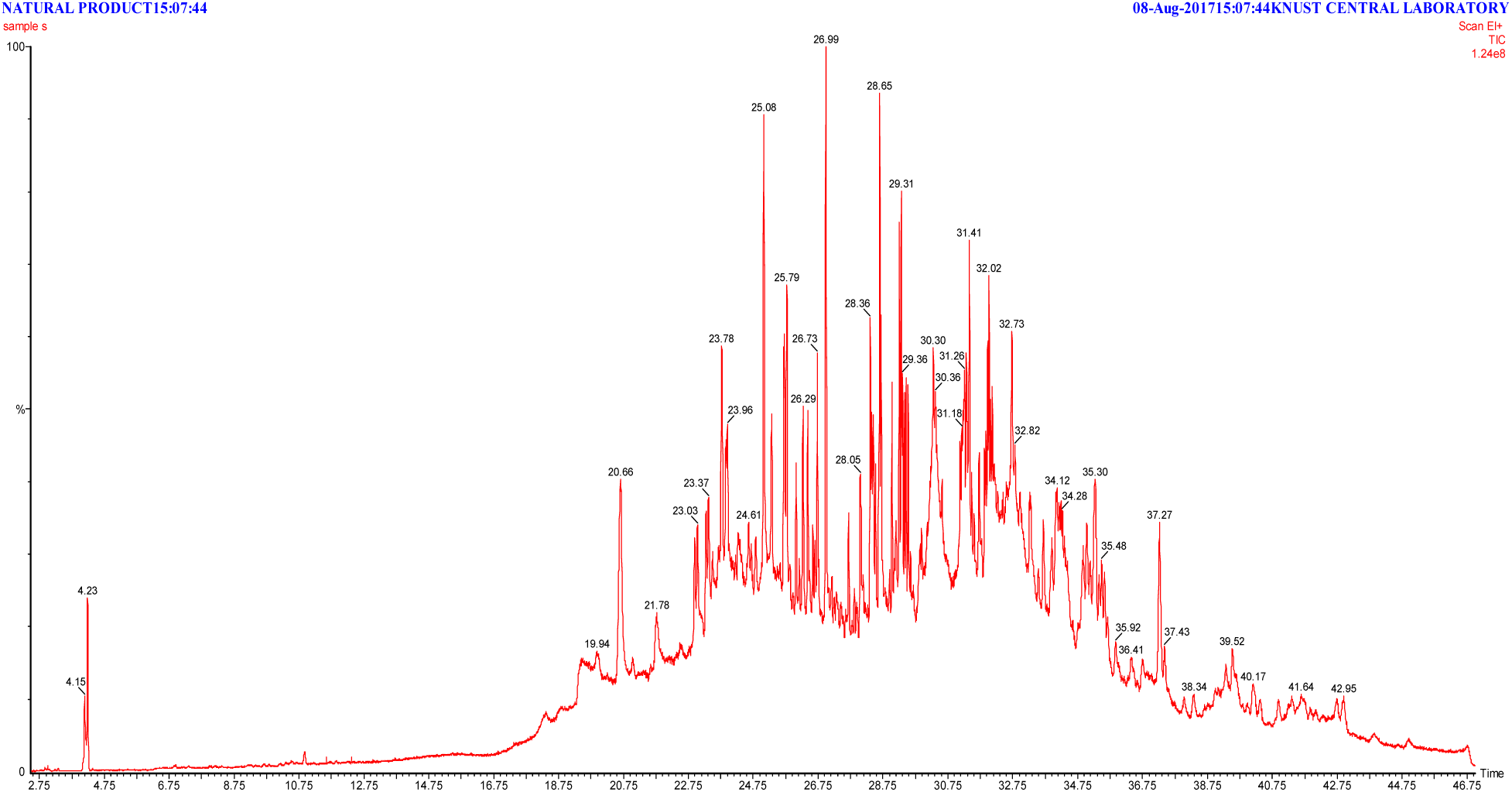
GC-MS chromatogram of fraction NKLS_6_2

**Figure 8.**
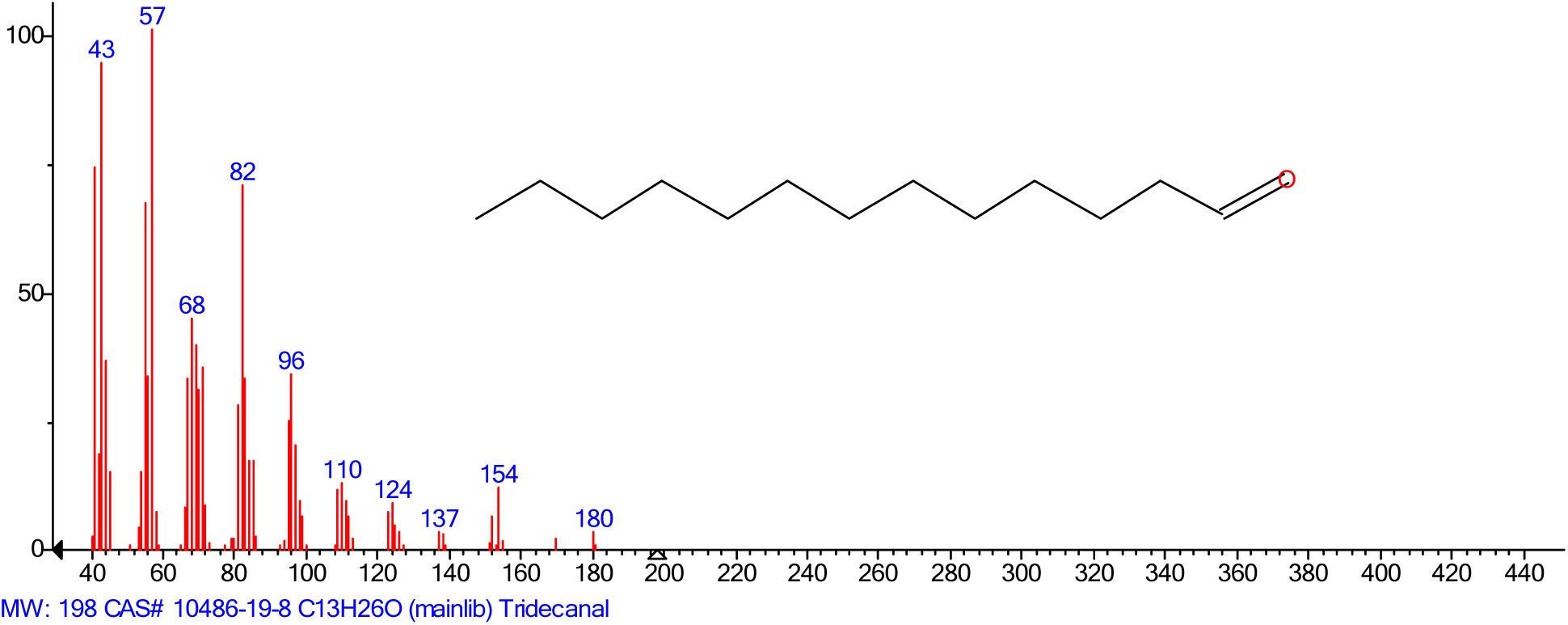
The chemical structure of the chromatogram Tridecanal (C_13_H_26_O)

**Figure 9.**
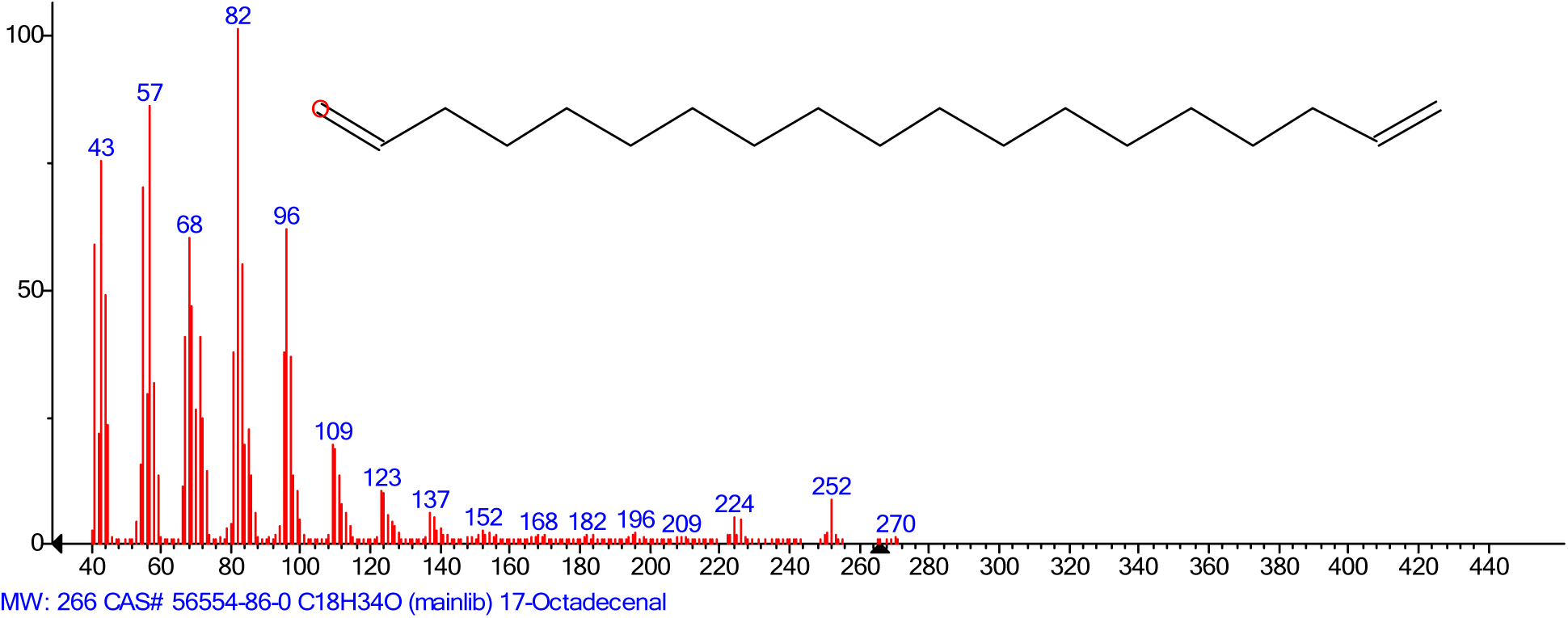
The chemical structure of the chromatogram 17-Octadecenal (C_18_H_34_O)

**Figure 10.**
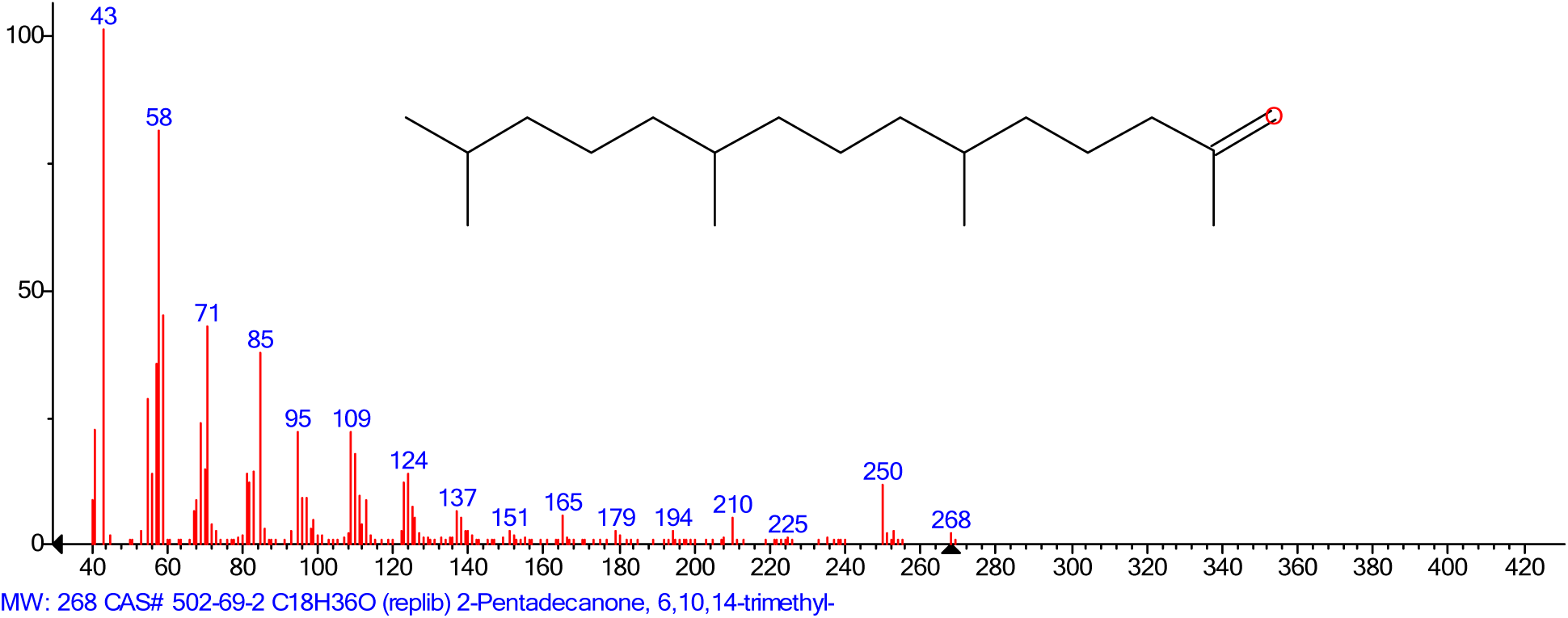
The chemical structure of the chromatogram Ethanol-2-(9-octadecenyloxy)-, (Z)

**Figure 11.**
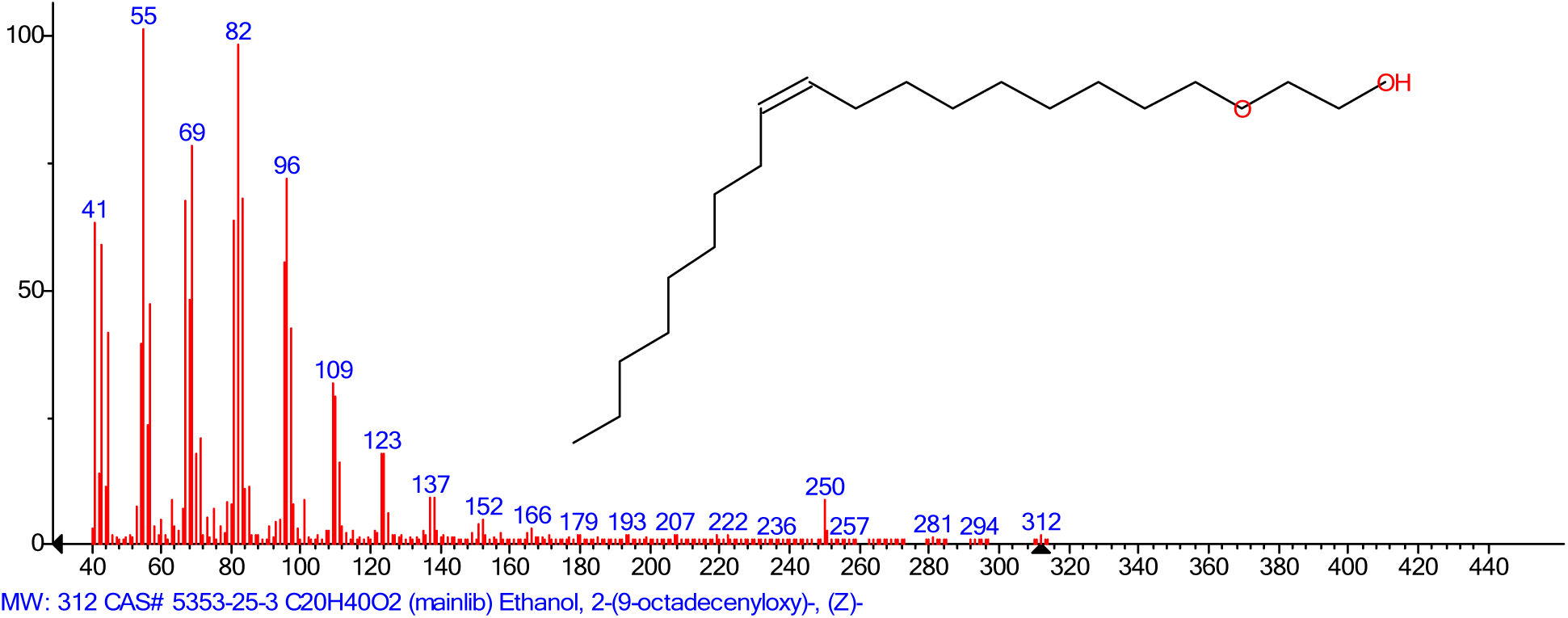
The chemical structure of the chromatogram 2-Pentadecanone-6, 10, 14, trimethyl (C18H36O)

**Figure 12.**
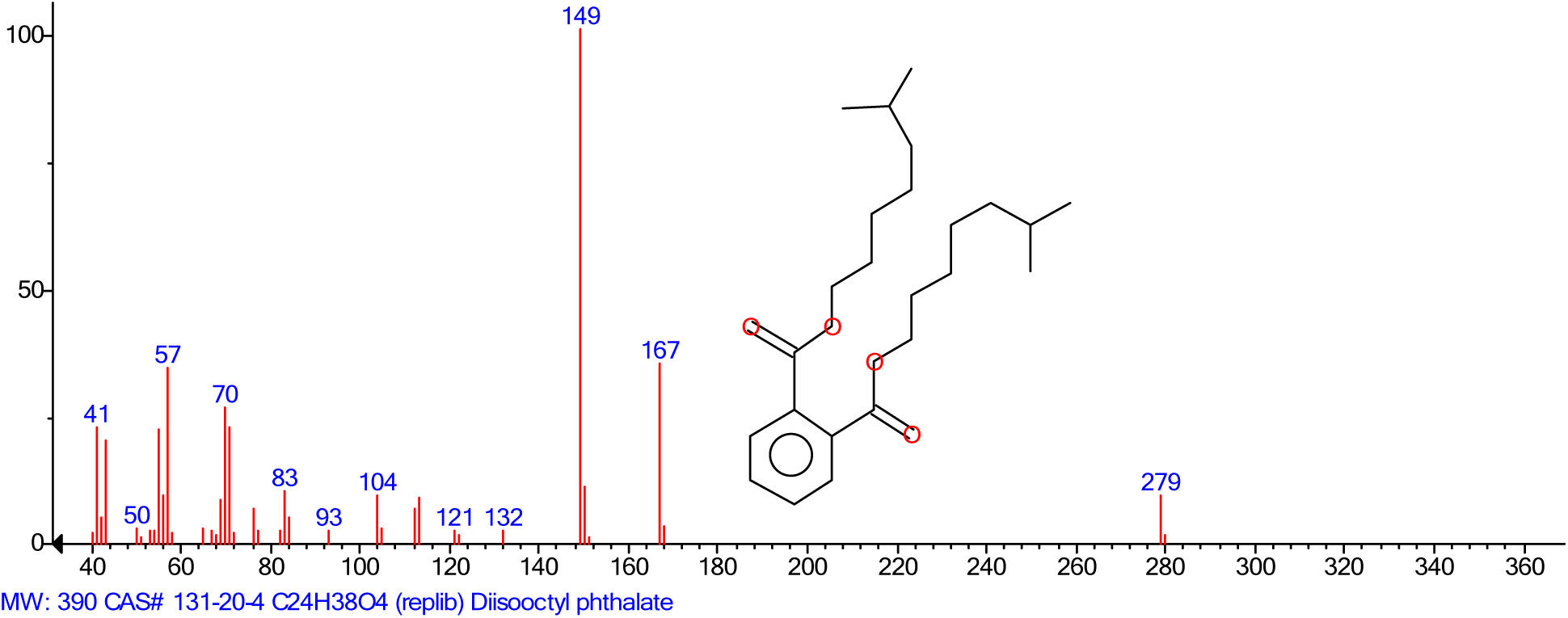
The chemical structure of the chromatogram Diisooctyl phthalate (C_24_H_38_O_4_)

**Table 7.** The results for the GC-MS of the fraction NKLS_6_2

| S/N | N. C. | M. F | R.T (min) | M. W | P. A (%) |
| --- | --- | --- | --- | --- | --- |
| (i) | Fatty aldehyde/ Fatty Acyls |  |  |  |  |
| 1 | Tridecanal | C<sub>13</sub>H<sub>26</sub>O | 20.66 | 198 | 2.05 |
| (ii) | Linoleyl Alcohol |  |  |  |  |
| 2 | 17-Octadecenal | C<sub>18</sub>H<sub>34</sub>O | 25.08 | 266 | 2.20 |
| (iii) | Polyethylene glycol oleyl ether or Alcoholic Compound |  |  |  |  |
| 3 | Ethanol-2-(9-octadecenyloxy)-,(Z) | C<sub>20</sub>H<sub>40</sub>O<sub>2</sub> | 26.99 | 312 | 1.67 |
| (iv) | Oxygenated Sesquiterpene |  |  |  |  |
| 4 | 2-Pentadecanone-6, 10, 14, trimethyl | C<sub>18</sub>H<sub>36</sub>O | 28.65 | 268 | 1.40 |
| (v) | Phthalic Acid |  |  |  |  |
| 5 | Diisooctyl Phthalate | C<sub>24</sub>H<sub>38</sub>O<sub>4</sub> | 37.29 | 390 | 1.26 |

S/N= Serial number, N.C= Name of the compound, MF= Molecular formula, R.T= Retention time, MW= Molecular weight, P.A= Peak area

## Discussion

Microorganisms are found in the environment such as air, water and soil ^3, 12^. This was ascertained when the Densu and Nsukwao rivers; lagoon and the sea samples produced 112 microorganisms after inoculation in a growth media. The medium was provided with protein, peptide or hydrocarbons which increase the microorganisms’ growth and production of antimicrobial metabolites as a result of protecting themselves from their prey as well as compete for nutrient ^13^. From the total of 112 microorganisms only 58 (51.79%) isolates yielded antimicrobial metabolites against at least two test organisms out of the eight test organisms used (Tables 1-3) and (Plate 1). Tawiah *et al.*^11^ isolated and screened 119 microorganisms and only 27 (22.69%) showed inhibitory activity against at least one of the test organisms. Also Kumar *et al.* ^’12^ isolated 78 marine actinobacteria and only 22 (28.21%) showed antibacterial and 12 (15.38%) antifungal activities. Water and sediment isolates (31) for actinomycetes were screened by Gebreyohannes *et al.* ^13^ and 13 (41.94%) isolates showed antibacterial activities against at least one of the test organisms.

The selected isolates, NKSEW_3_ and NKLS_6_ produce antimicrobial metabolites on the first day of incubation. On the 9^th^, 10^th^ and 11^th^ days of the incubation period, the isolates recorded the highest antimicrobial metabolites production per their zones of growth inhibitions against *Pseudomonas aeruginosa* (ATCC 27853), *Escherichia coli* (NCTC9002) and *Klebsiella pneumonia*e (clinical isolate) (Figure 1). These observations may suggest that the organisms started their antimicrobial metabolite production at their lag phase which continue into the exponential phase and became constant (stationary phase).

The aqueous extracts from the fermentation of the isolates NKSEW_3_ and NKLS_6_ demonstrated varied level of antimicrobial activity against the test organisms used (Table 4). The range of Minimum Inhibitory Concentrations (MIC) of the aqueous extracts ranged between 6.25-12.50 mg/mL for NKSEW_3_ and 3.13-12.50 mg/mL for NKLS_6_. This showed that some of the test organisms were more sensitive than the others (Table 5). The test microorganisms *E. coli*, *K. pneumoniae* and *P. aeruginosa* were killed at concentrations of 25.00, 12.50 and 25.00 mg/mL by NKSEW_3_ and 12.50, 6.25 and 25.00 mg/mL by NKLS_6_ respectively (Table 5). Ciprofloxacin was chosen because is a baseline drug that offers better absorption and a broad spectrum of in vitro activity against the test organisms used ^14^. Ciprofloxacin showed antimicrobial activity at lower concentration as compared to the extracts (Table 5).

Each microorganism had an optimum temperature for which it can grow, the selected isolates NKSEW_3_ and NKLS_6_ antimicrobial metabolites were subjected to temperature treatments before testing against *E. coli*, *K. pneumoniae* and *P. aeruginosa* using agar well diffusion. The isolates exhibited higher average zones of growth inhibition when incubated at 37 °C. At the temperature of 20 °C, 25 °C, 34 °C and 45 °C, the isolates NKSEW_3_ and NKLS_6_ have little effects for the production of antimicrobial metabolites (Figure 2). This work agrees with the studies conducted by Emelda and Vijayalakshmi ^15^ where antibiotic producing soil actinomycete were screened and antimicrobial activity was observed at 37 °C

Microorganisms grow well in their intracellular pH close to neutrality regardless of the pH outside their medium ^16^. The intracellular pH becomes stable when the gradient of the proton passing through the cytoplasmic membrane increases and forces the cells to charge resulting in cell lysis ^17^. Based on these, the effect of pH was assessed by varying the pH values in the fermentation medium. The isolates NKSEW_3_ and NKLS_6_ exhibited optimum antimicrobial activity at pH 8 when tested against *E. coli*, *K. pneumoniae* and *P. aeruginosa* (Figure 5). The work reported by Manjula *et al.* ^18^ was not different from this current study since they also reported that S*treptomyces* thrived best at pH 7-8 for antimicrobial production. Also report by Awais *et al*. ^19^ indicated that at pH 8 *Bacillus pumilus* produced metabolite with utmost inhibitory activity against *Micrococcus Luteus*. Work also done by Tawiah *et al*. ^11^ showed that strains identified as *Pseudomonas aeruginosa* produced metabolites with utmost activity at pH 7.

The growth and secondary metabolites biosynthesis by microorganisms are enhanced by a lot of factors for which carbon sources are part ^11^. These sources can be simple or complex carbohydrates ^11^. Based on these the isolates NKSEW_3_ and NKLS_6_ were cultivated in a medium containing glucose, xylose, glycerol, starch and galactose for their antimicrobial metabolites production. They were then tested against *E. coli*, *K. pneumoniae* and *P. aeruginosa*. The utmost production of antimicrobial metabolites of both isolates NKSEW_3_ and NKLS_6_ were recorded for the medium that contained glucose (Figure 3). The study reported by Gottschalk ^20^ was not different from this current study since he indicated that the quantity of glucose necessary for the growth of bacteria and antibiotic production differs according to the species of the bacteria. Tawiah *et al*. ^11^ corroborated the fact that glucose is a major source of carbon, an element essential for antimicrobial metabolite production.

Nitrogen sources also affect antimicrobial agents’ formation in microorganisms ^21^. Shapiro ^22^ indicated that the type of nitrogen source (organic or inorganic) plays a role in the synthesis of secondary metabolites. Farid *et al*. ^23^ reported that nitrates, nitriles and most ammonium salts are inorganic nitrogen sources which act by subduing the production of secondary metabolites and rather turn up increasing the biosynthesis of the cells. Madhava and Gnanamani ^24^ also revealed in their study concerning *Pseudomonas aeruginosa* MTCC 5210 that *Pseudomonas aeruginosa* MTCC 5210 produced utmost antimicrobial secondary metabolite using organic nitrogen sources rather than that of inorganic sources. These observations are consistent with the outcome of this work as ammonium sulphate and sodium nitrate produced utmost antimicrobial activity for isolates NKSEW_3_ and NKLS_6_ tested against *E. coli*, *K. pneumoniae* and *P. aeruginosa* (Figure 4).

Morphological and biochemical tests of the isolates NKSEW_3_ and NKLS_6_ conducted showed that both isolates were gram negative rod and they grew on cetrimide agar but not on MacConkey agar, mannitol Salt agar, bismuth sulphite agar. Also, they did not produce citrate, hydrogen sulphide, indole, nitrate but they produced acids from glucose and lactose. The study conducted by Tawiah *et al.*^11^ on characterization of selected isolates with antimicrobial activity gave similar results such that the microscopic examination of MAI 2 and MAI 3 disclosed similar features. As much as literature can ascertain from the observation and characteristics compared, no conclusion could be drawn on identity of organisms that absolutely matched the isolates NKSEW_3_ and NKLS_6._ Further identification was performed and the isolates were subcultured; identified using chromagar and biochemical tests. The results were compared with standard chart and the isolates were identified to be *pseudomonas species*. This result is not different from work reported by Samra *et al.* ^25^ where they used chromagar to identify different types of bacteria.

In order to purify the extracts, column chromatography was employed and fractions NKSEW_3_1and NKLS_6_2 were obtained tested for their antimicrobial activity. The fraction NKLS_6_2 exhibited high zone of growth inhibition against *E. coli*, *K. pneumoniae* and *P. aeruginosa* than NKSEW_3_1 (Table 6). This results support findings reported by Kyeremeh *el al.* ^26^ on fractions containing micromonospora for which only two showed antimicrobial activity. The fraction NKLS_6_2 was selected for HPLC analysis and the result revealed four peaks (Figure 6). The analysis of GC-MS of NKLS_6_2 revealed the presence of five compounds including; Tridecanal (C_13_H_26_O) a fatty aldehyde, an oxygenated sesquiterpene, 2-pentadecanone, 6, 10, 14-trimethyl (C_18_H_36_O), ethanol, 2-(9-octadecenyloxy)-, (Z) (C_20_H_40_O_2_), 17-octadecanal (C_18_H_34_O) and diisooctyl phthalate (C_24_H_38_O_4_) (Table 7). The ethanol, 2-(9-octadecenyloxy)-, (Z) (C_20_H_40_O_2_) has been reported to play an important role in antibacterial potential ^27^ whiles diisooctyl phthalate (C_24_H_38_O_4_) has been reported to be potential candidate for antimicrobial agent ^27, 28^. This work is not different from the study by Tulika, and Mala ^28^ that reported on GC-MS for the analysis of *E. crassipes* leaves which revealed the presence of 11 major compounds.

## 5.2 Conclusion

From the total of 112 isolated and screened microorganisms, 58 showed antimicrobial property against at least two of pathogenic organisms out of the eight using the agar well diffusion method. The isolates NKSEW_3_ and NKLS_6_ showed zone of growth inhibition ranged between 15 - 20 mm. The isolates NKSEW_3_ and NKLS_6_ were identified to be *Pseudomonas* species using chromagar and biochemical tests (oxidase test) to confirm. The temperature and optimum pH at which the utmost activity was recorded for both isolates (NKSEW_3_ and NKLS_6_) were 37 °C and 8, respectively. Glucose was the source of carbon that produced the utmost antimicrobial activity for both isolates NKSEW_3_ and NKLS_6_ while nitrogen sources were ammonium sulphate for NKSEW_3_ and sodium nitrate for NKLS_6_. The extracts of isolates NKSEW_3_ and NKLS_6_ showed MIC between 6.25-12.50 mg/mL for NKSEW_3_ and 3.13-12.50 mg/mL for NKLS_6_ and MBC between 12.5 – 25.00 mg/mL for NKSEW_3_ and 6.25 - 25.00 mg/mL for NKLS_6_ against *E. coli*, *K. pneumoniae* and *P. aeruginosa*. Column chromatography was used to fractionate the extracts and produced fractions NKSEW_3_1 and NKLS_6_2. The antimicrobial activity of the fraction NKSEW_3_1 and NKLS_6_2 showed that the fraction NKLS_6_2 exhibited higher zones of growth inhibition than NKSEW_3_1 against *E. coli*, *K. pneumoniae* and *P. aeruginosa* at a concentration of 100 mg/mL. The HPLC analysis of fraction NKLS_6_2 revealed four peaks while GC-MS revealed five compounds which including; Tridecanal, 17-octadecanal, ethanol, 2-(9-octadecenyloxy)-, (Z), 2-pentadecanone, 6, 10, 14-trimethyl and diisooctyl phthalate.

## Consent for Publication

Not applicable

## Data Availability

The data used to support the findings of this study are included within the article.

## Authors’ Contributions

Daniel Amiteye carried out the study and drafted the manuscript. Stephen Yao Gbedema, Vivian Etsiapa Boamah, Nicholas Tete Kwaku Dzifa Dayie, Francis Adu and Marcel Tunkumgnen Bayor designed the experiments, reviewed the manuscript, and supervised the work. All authors read and approved the final manuscript.

## Acknowledgement

The authors wish to express their gratitude to the Laboratory staff at the Department of Pharmaceutics, Faculty Pharmacy and Pharmaceutical Sciences, Kwame Nkrumah University of Science and Technology, Kumasi, Ghana for their technical assistance. Also, our great thanks go to Dr. Isaac Nortey Darko, University of Toronto, Canada for his assistance.

## Supplementary Materials

Effect of period of incubation on antibiotic metabolite production of the isolates, Effect of temperature on antimicrobial metabolite production of the isolates, Effect of pH on antibiotic metabolite production of the isolates, Effect of carbon sources on antibiotic metabolite production of the isolates, Effect of Nitrogen sources on antibiotic metabolite production of the isolates, Antimicrobial activity of the extracts, Gram reaction of isolates NKSEW_3_ and NKLS_6_ and Identification of isolate NKLS_6_ as *Pseudomonas species* using chromo agar.

## REFERENCES

1. Lerner, B. W., Lerner, K. L., editors, (2008). Infectious Diseases: Gale Group; Canada. Pp. 1 – 17, 50, 138 – 141 and 175 – 181.

2. Singh, A. P., Singh, R. B. and Mishra, S. (2012). Microbial and biochemical aspects of antibiotic producing microorganisms from soil samples of certain industrial area of India – an overview. The Open Nutraceuticals Journal. 5: Pp. 107 – 112.

3. Sarkar, P., Bhagavatula S., Suneetha, V., (2014). A brief research study on novel antibiotic producing isolate from VIT Lake, Vellore, Tamil Nadu. Journal of Applied Pharmaceutical Science. 4(1): 61 – 65.

4. Nike, A. R., Hassan, S. A., Ajijolakewu, M., Bosede. A. F. (2013). Soil screening for antibiotic – producing microorganisms. Advances in Environmental Biology. 7(1): 7 – 11.

5. Hirsch, C. F., Christensen, D. L. (1983). Novel Method for Selective Isolation of Actinomycetes. Applied and Environmental Microbiology. 925 – 929.

6. Sharma, M. (2014). Actinomycetes: source, identification, and their applications. Int. Journal Curr. Microbiol. App. Sci. 3(2):801 – 832.

7. Abdulkadir, M., Waliyu, S. (2012). Screening and Isolation of the Soil Bacteria for Ability to Produce Antibiotics. European Journal of Applied Sciences. 4 (5): 211 – 215.

8. Brumfitt, W., Hamilton-Miller, J.M. (1988). The changing face of chemotherapy. Postgrad Med Journal. 64(753): 552–558.

9. Mincer, T.J., Jensen, P.R., Kauffman, C.A., Fenical, W. (2002). Widespread and persistent populations of a major new marine actinomycete taxon in ocean sediments. Journal Appl Environ Microbiol, 68(10):5005–5011.

10. Radajewski, S., Webster, G., Reay, D.S., Morris, S.A., Ineson, P., Nedwell, D.B., Prosser, J.I., Murrell, J.C. (2002). Identification of active methylotroph populations in an acidic forest soil by stable isotope probing. Journal of Microbiol, 148:2331–2342.

11. Tawiah, A. A., Gbedema, S. Y., Adu, F., Boamah, V. E., Annan, K. (2012). Antibiotic producing microorganisms from River Wiwi, Lake Bosomtwe and the Gulf of Guinea at Doakor Sea Beach, Ghana. BMC Microbiology. 12: 234 – 242.

12. Kumar, K. S., Haritha R., Mohan J. Y. S. Y. V., Ramana T. (2011). Screening of marine Actinobacteria for antimicrobial compounds. Research Journal of Microbiology. 6(4): 385 – 393.

13. Gebreyohannes, G., Moges, F., Sahile, S. and Raja, N. (2013). Isolation and characterization of potential antibiotic producing actinomycetes from water and 82 sediments of Lake Tana, Ethiopia. Asian Pacific Journal of Tropical Biomedicine. 3(6): 426 – 435.

14. Neu, H. C., Percival, A., Lode, H. (1987). Ciprofloxacin: a major advance in quinolone chemotherapy. American Journal of Medicine. 82: 1–404.

15. Emelda, J., Vijayalakshmi, N. (2012). Isolation and Screening of Antibiotic Producing Soil Actinomycete for Antimicrobial Activity. Journal of Developmental Microbiology and Molecular Biology. 3(1): 47–54.

16. Garland, J. A. (1977). The dry deposition of sulphur dioxide to land and water surfaces. The Royal Society’s physical sciences research Journal. 2 (4) 102–114.

17. Riebeling, V., Thauer, R. K. & Jungermann, K. (1975). The internalalkaline pH gradient, sensitive to uncoupler and ATPase inhibitor, in growing Clostridium pasteurianum. Eur J Biochem. 55, 445–453.

18. Manjula, C., Rajaguru P., Muthuselvam, M. (2009). Screening for antibiotic sensitivity of free and immobilized Actinomycetes isolated from India. *Adv*. Biol. Res. 3: 84–88.

19. Awais, M., Qayyum, S., Pervez, A., Saleem, M. (2008). Effects of glucose, incubation period and pH on the production of peptide antibiotics by Bacillus pumilus. African Journal of microbiology research. 2(5):114–119.

20. Gottschalk, G. (1986). Bacterial Metabolism. Springer, Seoul. 52–60.

21. Merrick, M. J., and Edwards, R. A. (1995). Nitrogen control in bacteria. Microbiol. Rev. 59, 604–622.

22. Shapiro, S. (1989). Nitrogen assimilation in actinomycetes and the influence of nitrogennutrition on actinomycete secondary metabolism. In Regulation of Secondary Metabolism in Actinomycetes, Edited by S. Shapiro. Boca Raton, FL: CRC Press. 135–211.

23. Farid, M. A., Enshasy, H. A. E., Diwany A. I. E., Sayed, A. E. E. (2000). Optimization of the cultivation medium for natamycin production by *Streptomyces natalensis*. J. Basic Microbiol. 3: 157–166.

24. Madhava, C., Gnanamani, A. (2010). Condition Stabilization for *Pseudomonas aeruginosa* MTCC 5210 to Yield High Titers of Extra Cellular Antimicrobial Secondary Metabolite using Response Surface Methodology. Current. 3(4):197–213.

25. Samra, Z., Heifetz, M., Talmor, T., Bain, E., Bahar, J. (2002). Evaluation of use of a new chromogenic agar in detection of urinary tract pathogens. Journal of Clinical microbial. 36; 990–994.

26. Kyeremeh, K., Acquah, K. S., Sazak, A, Houssen, W., Tabudravu, Jioji, Deng, H., Jaspars M. (2014). Butremycin, the 3-Hydroxyl Derivative of Ikarugamycin and a Protonated Aromatic Tautomer of 5′-Methylthioinosine from a Ghanaian *Micromonospora sp*. K310. Mar. Drugs. 12: 999 – 1012.

27. Melo, I. S. Santos, S. N., Rosa, L. H., Parma, M. M., Silva, L. J., Queiroz, S. C. and Pellizari, V. H. (2014). Extremophiles 18(1), 15–23

28. Tulika, T., Mala, A. (2017). Phytochemical screening and GC-MS analysis of bioactive constituents in the ethanolic extract of *Pistiastratiotes* L. and *Eichhornia crassipes* (Mart.) solms. Journal of Pharmacognosy and phytochemistry. 6(1): 195–206.

